# Melatonin and GABA monotherapies are as efficacious as combination therapy in managing T1D and T2D: *In-vivo* studies on experimental diabetic models

**DOI:** 10.1101/2025.03.17.639558

**Authors:** Nishant Parmar, Roma Patel, Nirali Rathwa, Mitesh Dwivedi, Rasheedunnisa Begum, AV Ramachandran

**Author notes:** Corresponding Author: AV Ramachandran, Division of Life Science, School of Sciences, Navrachana University, Vadodara-391410, Gujarat, India. **Abbreviations:** T1D: Type 1 Diabetes; T2D: Type 2 Diabetes; M: Melatonin; GABA/G: γ-aminobutyric acid; FBG: Fasting Blood Glucose; STZ: Streptozotocin; HFD: High Fat Diet; TC: Total Cholesterol; TG: Triglyceride: HDL: High Density Lipoprotein; LDL: Low Density Lipoprotein; IPGTT: Intraperitoneal Glucose Tolerance Test; IPITT: Intraperitoneal Insulin tolerance Test; MTNR1A/MT1: Melatonin Receptor 1A; MTNR1B/MT2: Melatonin Receptor 1B; GABA_A_R: GABA receptor A; GABA_B_R: GABA receptor B; PDX1: Pancreatic and Duodenal Homeobox 1; ARX1: Aristaless-related homeobox-encoding gene; PAX4: Paired Box 4; NGN3: Neurogenin 3; AIF: Apoptosis Inducing Factor; BrdU: 5-bromo-2’-deoxyuridine; DAPI: 4′,6-diamidino-2-phenylindole; TUNEL: Terminal deoxynucleotidyl transferase dUTP nick end labeling; GCK: Glucokinase; FBPase: Fructose-1,6-Bisphosphatase; PEPCK: Phosphoenolpyruvate carboxykinase; GP: Glycogen phosphorylase; GS: Glycogen synthase; GLUT2: Glucose transporter 2; GLUT4: Glucose transporter 4; SIRT1: Sirtuin 1 PGC1α: Peroxisome proliferator-activated receptor gamma coactivator 1-alpha; ACC1: Acetyl-CoA carboxylase 1; ATGL: Adipose triglyceride lipase; IR1B: Insulin Receptor B; IRS: Insulin receptor substrate.

## Abstract

**Background:** Loss of functional β-cell mass is a critical factor in the ontogenesis of type 1 (T1D) and type 2 diabetes (T2D) marked by chronic hyperglycemia. Melatonin (M) and γ-aminobutyric acid (GABA/G) have immunoregulatory properties and have shown potential to increase β-cell mass in experimental diabetic models. Thus, we aimed to investigate the therapeutic potential of melatonin combined with GABA on β-cell regeneration in streptozotocin (STZ)-induced T1D and on a high-fat diet (HFD)-induced T2D mouse models.

**Methods:** The BALB/c mice were injected with 50 mg/kg body weight STZ on five consecutive days to induce T1D and were subjected to six weeks of monotherapy with melatonin or GABA and combination therapy (M+G). HFD was fed to C57BL/6J mice for 30 weeks to induce insulin resistance and T2D, followed by six weeks of mono or combination therapy exposure.

**Results:** Our studies on monotherapies and combination therapy reduced fasting blood glucose levels, increased plasma insulin levels and glucose tolerance, and promoted β-cell proliferation and transdifferentiation, with a simultaneous decrease in β-cell apoptosis in T1D mice. It also improved metabolic parameters and glycolipid metabolism in the liver and adipose tissue, respectively, and increased mitochondrial function in the skeletal muscle. Moreover, it restored peripheral insulin sensitivity and β-cell mass in T2D mice.

**Conclusion:** Our studies suggest that monotherapies are as effective as combination therapy since melatonin and GABA sub-additively ameliorates T1D and T2D manifestations.

## 1. Introduction

Type 1 diabetes (T1D) is an autoimmune disease characterized by progressive loss of functional β-cell mass resulting from insulitis, leading to insulin deficiency. In contrast, type 2 diabetes (T2D) is a metabolic disorder manifested by insulin resistance in peripheral tissues due to an impaired insulin signalling cascade [1]. The pathogenesis of T2D is more complex, involving different degrees of β-cell failure and insulin resistance. Obesity-induced insulin resistance and secretion defects are the major risk factors for T2D [1,2]. Both T1D and T2D ultimately result in pancreatic β-cell loss and chronic hyperglycemia [3–5]. Therefore, developing therapies that prevent or even reverse the deterioration of β-cell function is essential. Monotherapies often fail in the long run due to other diabetes-related complications and the patient’s lifestyle. Hence, combination therapy is recommended for long-term glycemic control [6].

Changing lifestyle trends such as a tendency to nocturnality and intake of high caloric diets cause disturbance in the sleep/wake cycle and circadian rhythms, favouring diabetes [7]. Melatonin, a pineal hormone, has a role in circadian rhythm regulation, acts as an antioxidant [8] and anti-inflammatory agent [9], and is also functionally linked to glucose metabolism [10]. There is an association between melatonin and T2D based on the findings that insulin secretion is inversely proportional to plasma melatonin concentration [11]. Pinealectomy leads to loss of melatonin, reduced glucose transporter type 4 (GLUT4) levels, glucose intolerance, and insulin resistance as melatonin checks insulin secretion [12,13]. Further, reduced melatonin levels are observed in various T2D rodent models and T2D patients [14]. Our previous study has shown that melatonin can induce β-cell regeneration in streptozotocin (STZ)-induced diabetic mice and β-cell regeneration in human islets transplanted in mice [4]. Insulin secretion and β-cell survival have been reported to be improved in response to melatonin signalling by decreasing β-cell apoptosis and oxidative stress in human islets exposed to chronic hyperglycemia and in islets from patients with T2D [15]. Moreover, studies on rodents have suggested that melatonin administration reduces body fat and Glycated haemoglobin (HbA1c) levels and increases *GLUT4* expression and insulin sensitivity in peripheral tissues in a diet-induced obese T2D mouse model [16].

Gamma (γ) - aminobutyric acid (GABA), initially identified as an inhibitory neurotransmitter and produced by pancreatic β-cells in large quantities, has emerged as a new anti-diabetic dietary supplement [17,18]. GABA has astonishing physiological effects in diabetes mellitus (DM) due to its anti-diabetic, antioxidant, anti-inflammatory, and immunomodulatory properties [18,19]. In an islet, GABA acts via GABA_A_R on the β-cells, enhancing insulin secretion through membrane depolarization [20]. Therefore, GABA is vital in regulating islet cell function and glucose homeostasis. Our previous study and other studies have shown that GABA-treated diabetic mice display higher plasma insulin and reduced glucagon levels, normalized glycemic control, and improved metabolic state in diabetic mice [5,21]. It stimulates β-cell replication, protects β-cells against apoptosis, attenuates insulitis, regulates the islet-cell function and glucose homeostasis, and suppresses detrimental immune reactions [5,21–25]. Recently, our group and others have shown the therapeutic effects of melatonin and GABA as monotherapies and in combination with other drugs (i.e. DPP-IV inhibitors, GLP-1R agonists) in diabetic models [4,5,22,26]. Hence, we aimed to assess the therapeutic potential of a combination of melatonin and GABA on diabetic manifestations in the STZ-induced T1D and HFD-induced T2D mouse models.

## 2. Materials and Methods

### 2.1 Animals

Forty male BALB/c mice (7-8 weeks old), bred in our animal vivarium, and forty young (6-7 weeks old) C57BL/6J male mice procured from ACTREC, Mumbai, were used for the experiment. The animals were maintained on a 12-hour light-dark cycle starting at 7.00 AM. These animals had free access to standard chow/HFD diet (Keval Sales Corporation, Vadodara, India) and water. All the experimental procedures were conducted as per the Purpose of Control and Supervision of Experiments on Animals (CPCSEA) guidelines and were approved by the Institutional Animal Ethical Committee (IAEC) (MSU/BC/02/2019; MSU/BC/09/2019).

### 2.2 Induction of T1D and treatment

For T1D induction, 32 BALB/c mice were given five consecutive intraperitoneal (i.p.) injections of 50 mg/kg body weight (BW) STZ (MP Biomedicals, Santa Ana, CA, USA), freshly dissolved in 0.1 M cold sodium citrate buffer (pH 4.5). Eight mice were kept separately as a non-diabetic control group. Diabetes was confirmed two weeks later in mice having fasting blood glucose (FBG) >350mg/dL. FBG was measured in mice after 6 hours of fasting by tail snipping using a Glucometer (TRUEresult, NIPRO Diagnostics, Pune, MH, India). The diabetic mice were then randomly assigned to four groups: i. Diabetic Control (DC) ii. Melatonin (M) treated iii. GABA (G) treated, and iv. M+G treated. Melatonin (Sigma, St. Louis, MO, USA) was administered between 6 PM and 7 PM daily at 0.5 mg/kg BW i.p. in 0.9% saline [4]. GABA (Sigma–Aldrich, United States) was given at 18 mg/day [22] by oral gavage. Control and DC groups received 0.9% saline and water as vehicle control for M and G. The treatment was given for six weeks along with bromodeoxyuridine (BrdU) (MP Biomedicals, Santa Ana, CA, USA) on alternative days at 100 mg/kg BW, i.p. dose. FBG levels and BW were measured twice a week (Fig. S1).

### 2.3 Induction of T2D and Treatment

For T2D induction, 32 C57BL/6J mice were fed a high-fat diet (HFD), and eight control mice were fed a standard chow diet (NCD) for 30 weeks. T2D was confirmed 30 weeks later in mice having FBG >200 mg/dL. HFD-fed mice were randomly assigned into four groups: i. HFD ii. Melatonin (M) treated iii. GABA (G) treated, and iv. M+G treated. Melatonin (Sigma, St. Louis, MO, USA) was administered between 6 PM and 7 PM daily at 10 mg/kg BW i.p., dissolved in 0.1% ethanol and 0.9% saline [4]. GABA (Sigma–Aldrich, United States) was given at 10 mg/day [22] by oral gavage. NCD and HFD groups received 0.1% ethanol + 0.9% saline and water as vehicle control for M and G, respectively. The treatment was given for six weeks. FBG levels and BW were measured weekly, along with food and water intake (Fig. S2).

### 2.4 Intraperitoneal Glucose Tolerance Test (IPGTT) and Intraperitoneal Insulin Tolerance Test (IPITT)

Glucose tolerance and insulin sensitivity were evaluated by IPGTT and IPITT, respectively, at the end of treatment. For IPGTT and IPITT, all mice were fasted for 6 hours and injected with glucose (2 g/kg BW i.p.) and insulin (0.75 U/kg BW), respectively. Blood glucose levels were measured immediately at 0, 15, 30, 60, 90, and 120 minutes by tail snipping method using a Glucometer. The total area under the curve (AUC) was evaluated.

### 2.5 Plasma Parameters

1 ml blood was collected into K3 EDTA tubes using the orbital sinus method for the biochemical assays after 6 hours of fasting before sacrificing mice. Additionally, tissues were harvested, which are further described in depth below. Blood was centrifuged at 6000 g for 5 minutes, and plasma was separated and stored at −20°C for further analysis. Plasma insulin, leptin, and melatonin were measured by commercially available mouse insulin, mouse leptin (both from RayBiotech, GA, USA), and mouse melatonin (Elabscience, Houston, TX, USA) ELISA kits as per the manufacturer’s protocols. Lipid profile (triglycerides, total cholesterol, HDL-c) was measured by commercially available kits (Reckon Diagnostics Pvt. Ltd., Vadodara, GJ, India). Friedewald’s (1972) formula calculates low-density lipoprotein (LDL).

### 2.6 Gene Expression Analysis

Parts of the liver, skeletal muscle, and adipose tissue were stored in RNAlater™ Stabilization Solution (Thermo Fisher Scientific, USA) for gene expression analysis of key glucoregulatory genes and *glucose transporter 2* (*GLUT2*), mitochondrial biogenesis, and lipid metabolism, respectively. Total RNA was extracted by the Trizol method, as described previously [27]. The expression of targeted genes and *GAPDH* transcripts were monitored by LightCycler®480 Real-time PCR (Roche Diagnostics GmbH, Germany) using gene-specific primers (Eurofins, India), as shown in Table S1. The expression of the *GAPDH* gene was used as a reference. Real-time PCR was performed as described previously by determining 2^-ΔΔCt^ [27].

### 2.7 Droplet Digital PCR (ddPCR)

Gene expression of *MTNR1B* and *GLUT4* in mouse adipose tissue was monitored by ddPCR using EvaGreen dye (Bio-Rad, Hercules, CA, USA). Each ddPCR reaction contained 20 µl of the master mix, including 10 µl of Eva Green supermix, 0.5 µl (2.5 mM) forward and reverse primer each, 1 µl cDNA (50 ng), and 8 µl nuclease-free water. This system was loaded into an oil well in a droplet generator cartridge with 70 µl of droplet generation oil. A QX200 TM Droplet Generator was used to generate droplets loaded onto 96-well PCR plates. The plate was heat-sealed with foil and was run in a thermal cycler. The PCR conditions for the assay were: 95 °C for 5 min; 40 cycles of 95 °C for 30s and 60 °C (depending on annealing temperature) for 1 min; and three final steps at 4 °C for 5 min, 90 °C for 5 min, and a 4 °C for 30 min to boost dye stabilization. The PCR product was read in QX200TM Droplet Reader and was analyzed by QuantaSoft™ software. The results were plotted as Ch1Amplitute and copies/μl. Blue droplets show several positive droplets for their respective target genes, and grey droplets indicate negative droplets for the target genes. Non-template control was run for each assay. The details of the forward and reverse primers are shown in Table S1.

### 2.8 Glucoregulatory Enzyme Activity and Liver Glycogen Content

A portion of the liver tissue was harvested and snap-frozen with dry ice, then stored at −80 °C for enzyme activity assays. 50 mg tissue was homogenized, and tissue lysates were used for the activity assays. Enzyme assays for Glucokinase (GCK), Fructose-1,6-Bisphosphatase (FBPase), phosphoenolpyruvate carboxykinase (PEPCK), Glycogen phosphorylase (GP), and glycogen content were carried out by commercially available kits (BioVision, Milpitas, CA, USA) according to the manufacturer’s protocol.

### 2.9 Mitochondrial Oxygen Consumption Rate (OCR)

A portion of skeletal muscle was harvested and stored in a mitochondrial respiration buffer at −80 °C for OCR studies. Mitochondria were isolated from skeletal muscle using a mitochondria isolation kit (Thermo Scientific TM, Catalog no. 89801) using the manufacturer’s protocol. The isolated mitochondria were resuspended in 100 µl of mitochondria respiration buffer (110mM Sucrose, 0.5mM EGTA, 70 mM KCl, 0.1% FFA free BSA, 20 mM HEPES, 3 mM MgCl2, and 10 mM KH2PO4, 20mM Taurine) (Butterick et al., 2016). OCR studies were conducted in the Oxytherm System (Hansatech Instruments Ltd., Pentney, UK). Outer membrane integrity of the isolated mitochondria was evaluated by impermeability to exogenous cytochrome c, which was constantly >95%. The activities of Respiratory chain complexes I-IV were monitored using 100 µl of substrates [100 mM Pyruvate & 800 mM Malate (complex I), 1 M Succinate (complex II), 10 mM α-glycerophosphate (complex III), and 0.8 M ascorbate (complex IV)] (Li and Graham, 2012) added to 90 µl of mitochondria suspension. Other respiration reagents used were 100 mM adenosine diphosphate (100 µl), 1 mM oligomycin (2 µl), 1 mM rotenone (1 µl), and 1 mM Antimycin (2.5 µl). Protein concentration was estimated using the Bradford method [28]. All chemicals were purchased from Sigma-Aldrich, USA. OCR was determined by measuring the amount of oxygen (nmol) consumed, divided by the time elapsed (min) and the amount of protein present in the assay [29]. Data represents the respiratory control ratio (RCR) of state 3/state 4 respiration.

### 2.10 Western Blot Analysis

The skeletal muscle was also harvested and stored in a lysis buffer containing protease inhibitor cocktail and phosphatase inhibitor cocktail 2 and 3 (both from Sigma, St. Louis, MO, USA) at −80 °C for western blot analysis of key proteins involved in the insulin signalling pathway. The tissue was homogenized in liquid nitrogen and Laemmli buffer (1M Tris HCl, 10%SDS, 20% glycerol, and 10% β-mercaptoethanol) containing 1 M urea (1:1). The homogenate was collected and sonicated twice in cold conditions and centrifuged to remove tissue/cell debris. After performing the TCA precipitation method, the protein concentration in the lysates was estimated using the Bradford assay [28]. 25-40 µg of the lysate were resolved on 8-10% SDS-PAGE, followed by electrophoretic transfer to the PVDF membrane. Membranes were blocked with 5% bovine serum albumin (Sisco Research Laboratories Pvt. Ltd., Mumbai, MH, India) or Blotting-Grade Blocker (Bio-Rad) in tris-buffered saline (pH 8.0) with 0.1% Tween-20 for 1 hour at room temperature. For immunoblot analysis, the membrane was incubated overnight with different targeted primary antibodies at 4°C. Membranes were then incubated with secondary antibodies at room temperature for 1 hour. The details of primary and secondary antibodies with dilutions are shown in Table S2. The membrane was visualized with the clarity western ECL substrate (Bio-Rad Laboratories, USA) in the ChemiDoc™ Touch Imaging System. Blots were analyzed using Image LabTM software (Bio-Rad Laboratories, USA).

### 2.11 Pancreatic Tissue Preparation and Immunohistochemistry-Immunofluorescence (IHC-IF)

The pancreas was fixed in 10% neutral buffered formalin for histological processing and paraffin embedding. 5µm sections were prepared from the paraffin-embedded blocks. Immunofluorescence staining was carried out to study β-cell proliferation (Insulin/Glucagon/BrdU), β-cell neogenesis (Insulin/NGN3/PDX1), α-to β-cell transdifferentiation (Insulin/ARX/PAX4), β-cell apoptosis (TUNEL assay kit; Thermo Fisher Scientific, MA, USA) and Insulin/apoptosis-inducing factor (AIF) staining, and β-cell mass & islet number (Insulin staining). The sections were deparaffinized in xylene and rehydrated in a series of graded ethanol (100%, 95%, 80%, and 70%). Antigen retrieval was performed using 1N HCL for 45 minutes at 37 °C. Sections were blocked in 5% donkey serum in PBST (PBS + 0.1% Tween 20) for 1 hour at room temperature, and antibodies were diluted as indicated in Table S3. The sections were incubated with primary antibody at 37°C in a humidified chamber for 2 hours. Tissue sections were washed with PBS and incubated with secondary antibodies at room temperature for 45 minutes in the dark. Tissue sections were washed with PBS and distilled water and were mounted with Slowfade Gold Antifade mountant with DAPI (Thermo Fisher Scientific, USA), and the coverslip was sealed with nail varnish. Stained sections were observed under a confocal laser scanning microscope (Zeiss LSM 780, Oberkochen, Germany) at 63X for β-cell regeneration and apoptosis. Image analysis was carried out using Zeiss Zen software. Results were expressed as the percentage of specified markers for β-cell regeneration and apoptosis [4]. Whole pancreatic tissue images were acquired on the tile-scan mode (with automatic image stitching) under a laser scanning microscope (Leica SP8 TCS, Mannheim, Germany) at 10X magnification for β-cell mass & islet number, and Image analysis was carried out in Image J software. Observations were made from three pancreatic sections per group from five different areas.

### 2.12 Statistical Analyses

Statistical data comparisons were performed by one-way analysis of variance (ANOVA) and multiple group comparisons by Tukey’s post hoc test in GraphPad Prism 6 (GraphPad Sofware, San Diego, CA, USA). The significance level was set as *p*<0.05. Results are expressed as mean±SEM.

## 3. Results

### 3.1. Effect of melatonin, GABA, and combination treatment on pancreatic β-cell regeneration in Streptozotocin (STZ) - induced T1D mouse model

#### 3.1.1 Assessment of Body Weight, Fasting Blood Glucose, and Glucose Tolerance

Metabolic profiling in mice suggests that the DC group showed a significant increase in FBG levels (*p*<0.001) with a reduction in plasma insulin levels (*p*<0.001) and glucose tolerance (*p*<0.001) as compared to the control group. The monotherapies and the combination therapy significantly reduced FBG levels (M, *p*<0.001; G, *p*<0.001; M+G, *p*<0.001) by increasing plasma insulin levels (M, *p*<0.01; G, *p*<0.001; M+G, *p*<0.001) with a consequent increase in glucose tolerance (M, p<0.001; G, p<0.001; M+G, p<0.001) as compared to DC group post-treatment.

Furthermore, final FBG levels in the drug-treated groups were also significantly reduced as compared to their initial levels (M, *p*<0.01; G, *p*<0.001; M+G, *p*<0.001). However, all the treated groups showed no significant change in body weight before and after treatment (Figure 1A-E).

**Figure 1:**
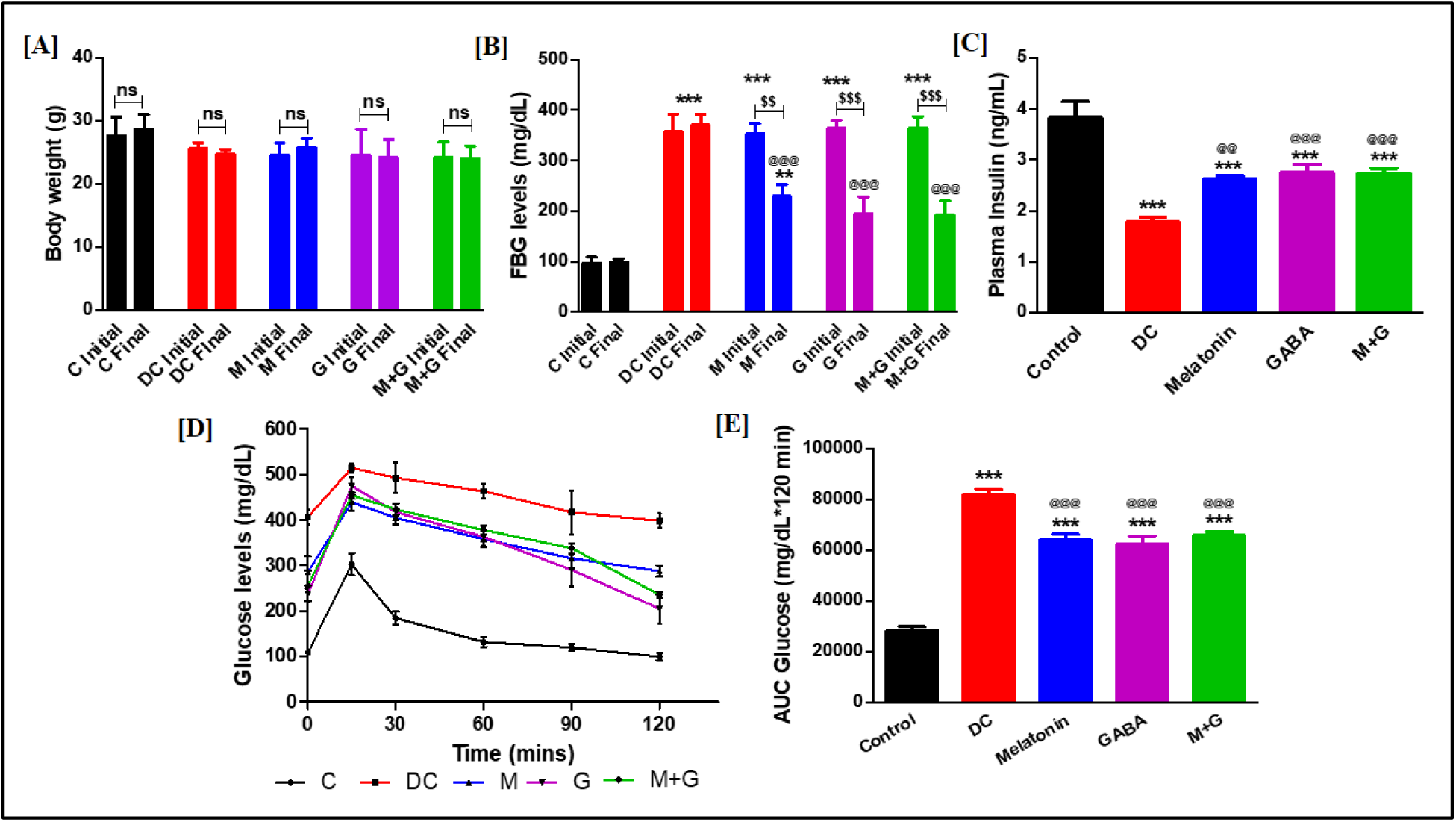
Assessment of Body Weight, Fasting Blood Glucose levels, and Glucose Tolerance. **[A] Body weight**. No significant change in body weight was observed before and after treatment in all the treated groups. **[B] Fasting blood glucose levels**. A significant increase in FBG levels was observed in the DC group compared to the control group. In contrast, a significant decrease in FBG levels was observed in all the drug-treated groups compared to their initial levels. Also, there was a significant reduction in the FBG levels in the final drug-treated groups compared to the final DC group. **[C] Plasma insulin levels**. A significant decrease in random plasma insulin levels was observed in the DC group compared to the control group. In contrast, the levels were significantly increased in all the drug-treated groups compared to the DC group. **[D] Glucose tolerance test**. At all times, a significant decrease in glucose clearance was observed in the DC group compared to the control group. Increased glucose clearance was observed in all the drug-treated groups compared to the DC group at all times. **[E] Blood glucose AUC0–120**. A significant decrease in glucose tolerance was observed in the DC group compared to the control group. In contrast, all the drug-treated groups showed a significant increase in glucose tolerance compared to DC. (\*\**p*<0.01, \*\*\**p*<0.001 vs. NCD; ^@@^*p*<0.01, ^@@@^*p*<0.001 vs. HFD; ns, *p>*0.05) (n=6/8 per group).

#### 3.1.2 Assessment of Pancreatic β-cell regeneration and Apoptosis

IHC analysis showed no change in β-cell proliferation and transdifferentiation in the DC group compared to the control group, as shown by BrdU^+^ insulin^+^ cells and PAX4^+^ ARX^+^ insulin^+^ cells, respectively. In comparison, it was significantly increased in all the drug-treated groups (proliferation: M, *p*<0.01; G, *p*<0.01, M+G, *p*<0.001; transdifferentiation: M, *p*<0.01; G, *p*<0.001, M+G, *p*<0.001). Moreover, a nucleo-cytoplasmic translocation of PAX4 was observed in DC and all the drug-treated groups. Intriguingly, the analysis revealed no β-cell neogenesis as NGN3 was found negative. However, we observed PDX1+ insulin+ cells in all the groups indicated by the nucleo-cytoplasmic translocation of PDX1. Additionally, caspase-independent β-cell apoptosis was not observed in any group, as shown by negative AIF translocation into the nucleus of insulin^+^ cells. However, β-cell apoptosis, as shown by TUNEL^+^ insulin^+^ cells, was significantly increased in the DC group (*p*<0.001) as compared to the control group (*p*<0.001), while it was reduced significantly in all the drug-treated groups (M, *p*<0.01; G, *p*<0.05, M+G, *p*<0.01) (Figure 2A-E).

**Figure 2:**
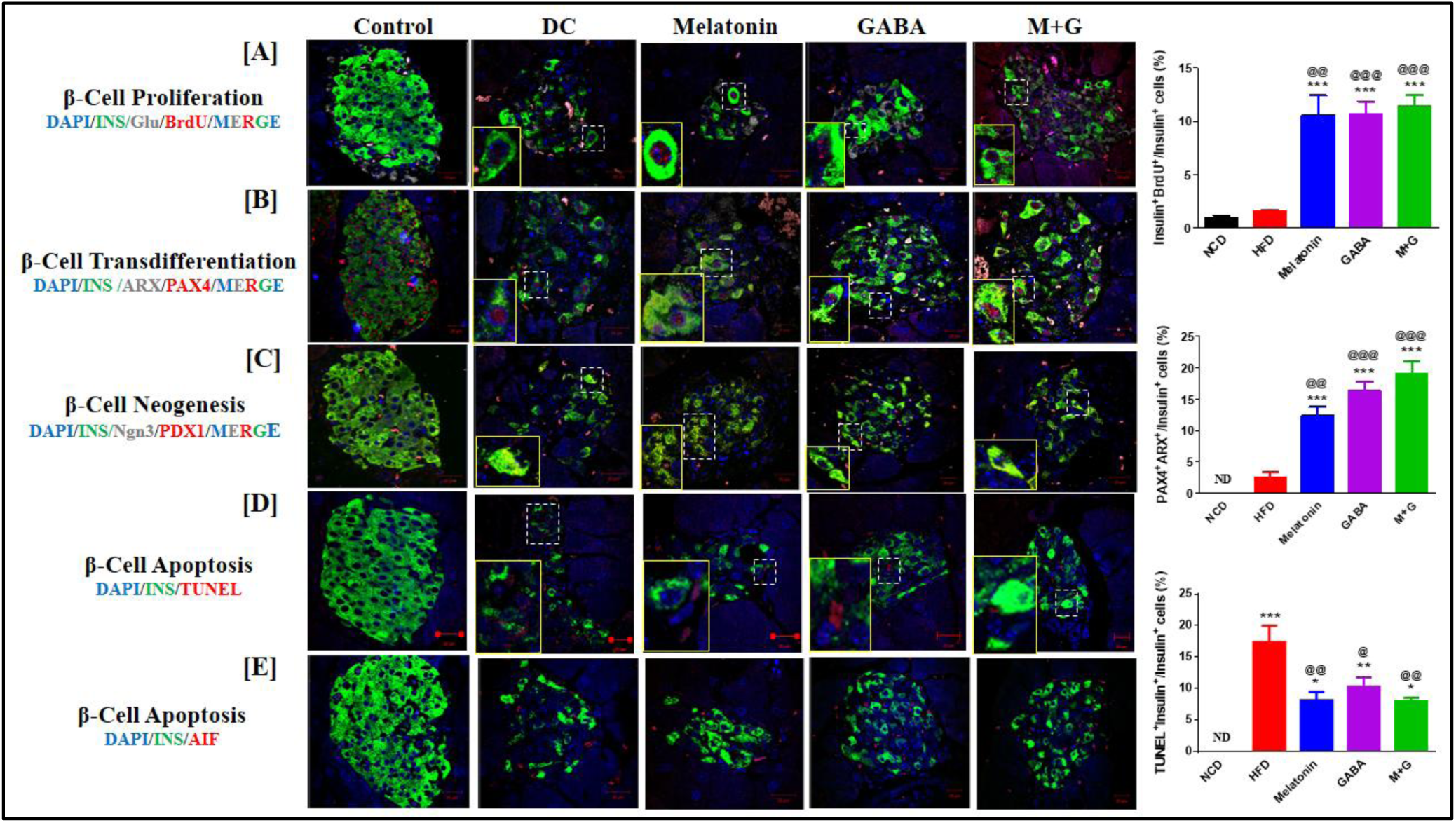
Assessment of Pancreatic β-cell regeneration and Apoptosis. **[A] β-Cell Proliferation.** No significant change in β-cell proliferation was observed in the DC group compared to the control group. All treated groups showed a significant increase in β-cell proliferation compared to the DC group. **[B] β-Cell transdifferentiation.** α-to β-cell transdifferentiation was observed in DC and all the drug-treated groups, while it was significantly increased in all the drug-treated groups compared to the DC group. Moreover, a nucleo-cytoplasmic translocation of PAX4 was also observed in DC and all the drug-treated groups. **[C] β-Cell neogenesis.** NGN3^+^ cells were not detected in DC or the drug-treated groups. However, PDX1^+^ cells were detected in all the groups with their nucleo-cytoplasmic translocation. **[D] β-Cell Apoptosis by AIF.** None of the groups showed AIF translocation to the nucleus in insulin^+^ cells. **[E] β-Cell Apoptosis by TUNEL.** β-cell apoptosis was significantly increased in the DC group compared to the control group and significantly reduced in all the treated groups compared to the DC group. Scale-20 µm. Magnification-63X. (\**p*<0.05, \*\**p*<0.01, \*\*\**p*<0.001 vs. NCD; ^@^*p*<0.05, ^@@^*p*<0.01, ^@@@^*p*<0.001 vs. HFD) (n=3/group).

### 3.2 Effect of melatonin, GABA, and combination treatment on High Fat Diet (HFD) - induced T2D mouse model

#### 3.2.1 Assessment of Metabolic Profile

Our results suggest that monotherapies (M, *p*<0.01; G, *p*<0.01) are as efficacious as combination therapy (p<0.001) in reducing FBG levels, increasing glucose tolerance (S, *p*<0.01; M, *p*<0.05; M+G, *p*<0.001) and insulin sensitivity (S, *p*>0.05; M, *p*>0.01; S+M, *p*<0.001). The BW of the HFD group was significantly increased compared to the NCD group (*p*<0.001). However, the treated groups did not significantly reduce the final BW compared to their initial BW (*p*>0.05). Moreover, an assessment of food and water intake revealed that there was a significant reduction in food intake (HFD, *p*<0.05; M, M+G; *p*<0.01), and no difference was observed in water intake (HFD, M, G, M+G; *p*>0.05) after six weeks of the drug treatment as compared to NCD group. Further, a notable change in water intake was observed before and after treatment in the M+G treated group (*p*<0.05) (Figure 3A-F).

**Figure 3:**
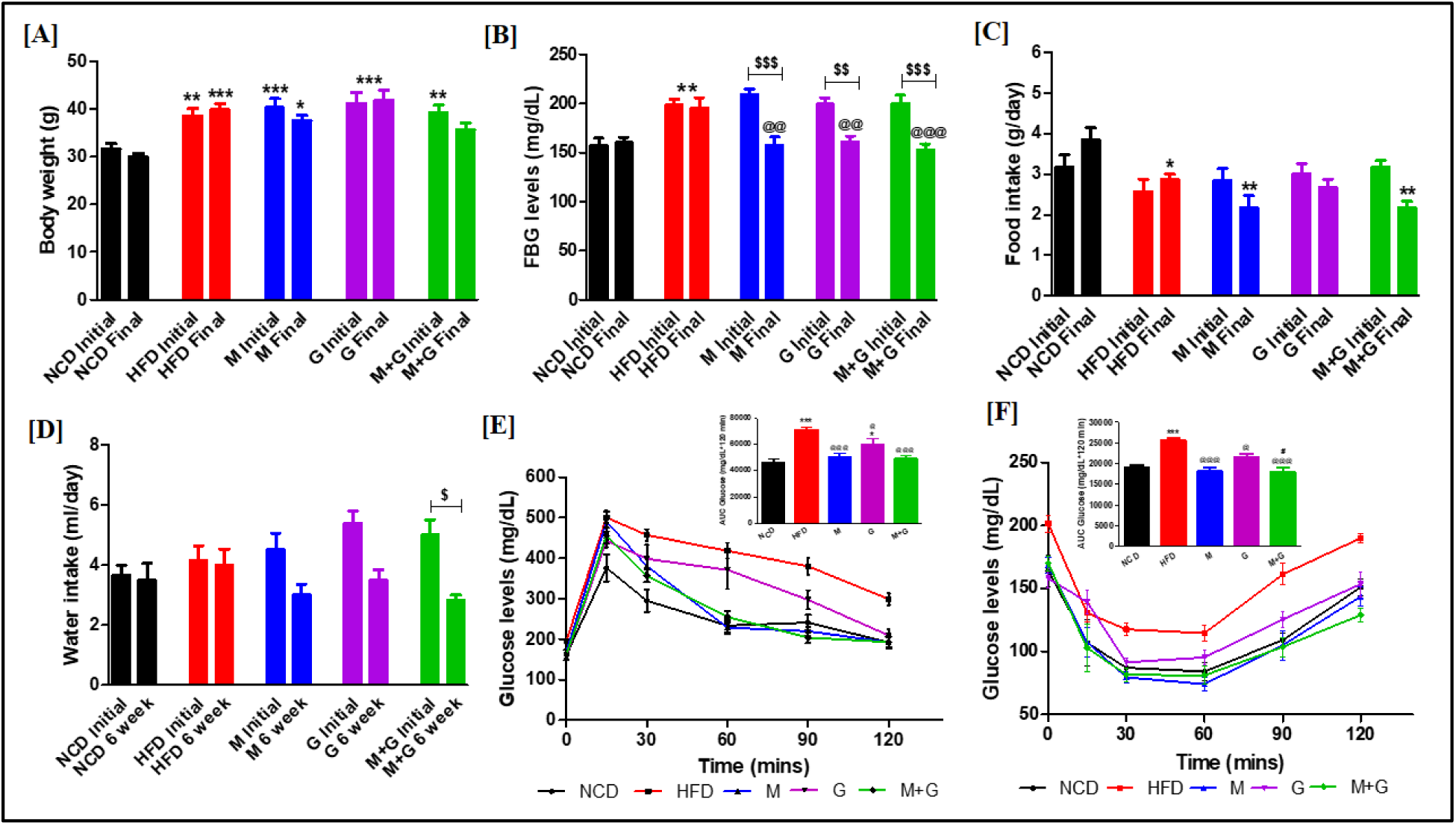
Bodyweight, blood glucose levels, food and water intake, IPGTT and IPITT. **[A] Bodyweight. [A] Bodyweight.** A significant increase in body weight was observed in HFD-fed mice compared to NCD. No significant difference was observed in the body weight six weeks post-drug treatment. **[B] Fasting blood glucose levels.** A significant increase in FBG levels was observed in HFD-fed mice and significantly reduced in all treated groups after six weeks post-drug treatment. **[C] Food intake.** No significant difference was observed in the food intake after six weeks post-drug treatment. HFD-fed and other drug-treated groups showed a reduction compared to NCD. [**D] Water intake.** A significant reduction in water intake was observed in M+G treated groups after six weeks post-drug treatment **[E] lPGTT and AUC glucose curve:** IPGTT. Increased glucose clearance was observed in all the treated groups compared to HFD-fed mice. **Blood glucose AUC0–120.** All the treated groups showed a significant increase in glucose tolerance compared to HFD-fed mice. **[F] lPITT and AUC glucose curve:** Increased glucose clearance was observed in all the treated groups compared to HFD-fed mice. **Blood glucose AUC0–120.** All the treated groups showed a significant increase in insulin sensitivity compared to HFD-fed mice. (\**p*<0.05, \*\**p*<0.01, \*\*\**p*<0.001 vs. NCD; ^@^*p*<0.05, ^@@^*p*<0.01, ^@@@^*p*<0.001 vs. HFD) (n=6-8/group).

#### 3.2.2 Assessment of Plasma Lipid Profile, Insulin, Leptin and Melatonin levels

Analysis of plasma lipid profile suggested that the levels of HDL were increased in the HFD group but were not significant, but TG, TC, and LDL were significantly increased in the HFD group (TG, *p*<0.001; TC, *p*<0.01; LDL, *p*<0.05) as compared to NCD indicating dyslipidemia. Lipid levels (TG, TC, and LDL) were restored in the monotherapies and combination-treated groups as compared to the HFD group (TG: M, G, M+G, *p*<0.001; TC: M, *p*<0.01; G, M+G, *p*<0.01; LDL; M, G, M+G, *p*<0.01). However, no changes in HDL levels were observed in any of the treated groups (p>0.05). Furthermore, Analysis of plasma insulin, leptin, and melatonin levels revealed that there was hyperinsulinemia and hyperleptinemia along with decreased melatonin levels in the HFD group as compared to NCD (*p*<0.001), and the levels were restored in the drug-treated groups (insulin: M, M+G, *p*<0.01; G, *p*<0.05; leptin: M, M+G, *p*<0.05; melatonin: M, M+G, *p*<0.001) as compared to HFD group (Figure 4A-G).

**Figure 4:**
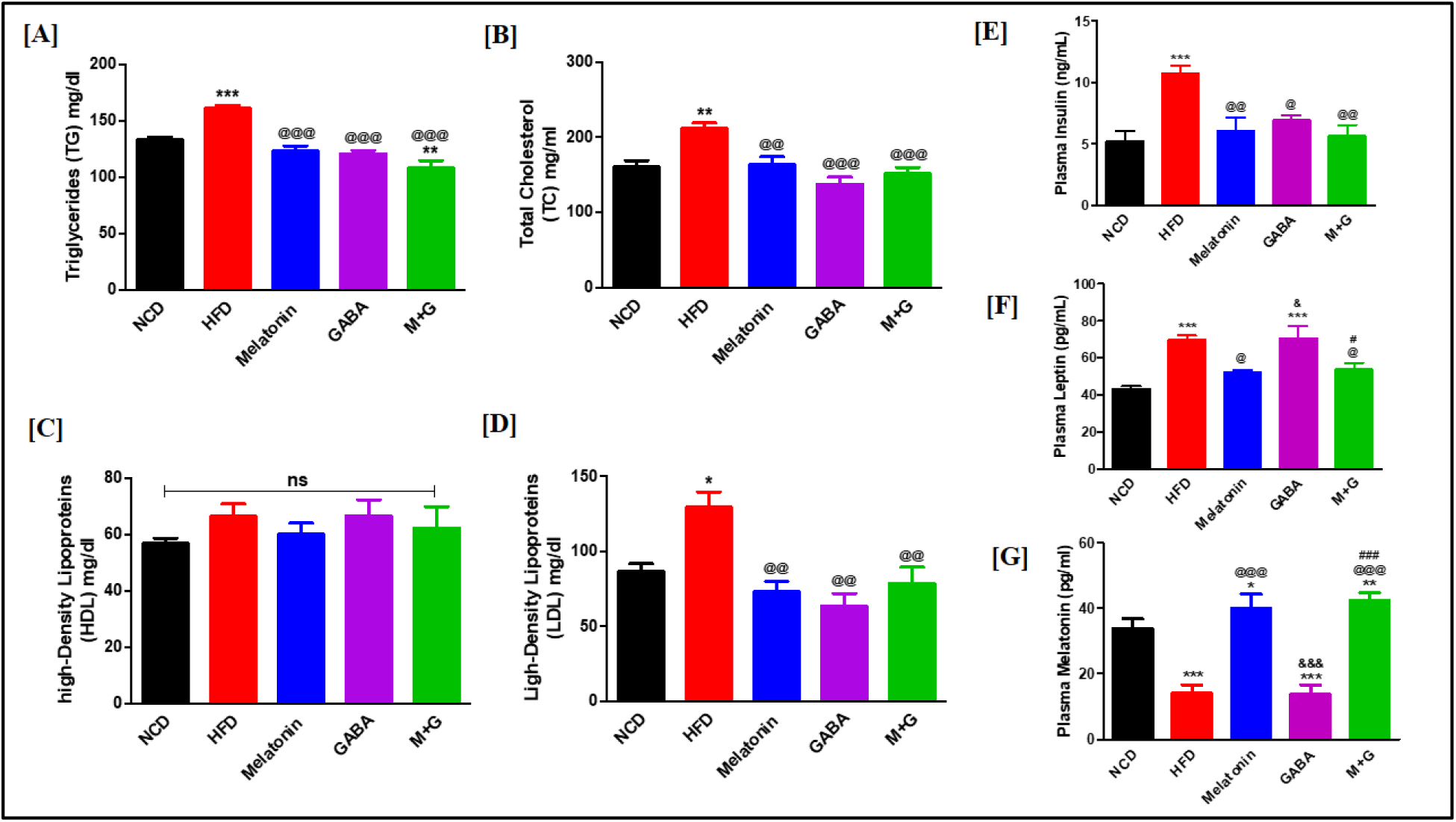
Plasma Lipid Profile, Insulin, Leptin, and Melatonin levels. A significant reduction of triglycerides **[A]**, total cholesterol **[B],** and LDL **[D]** was observed in all the treated groups as compared to HFD-fed mice. In comparison, HDL levels were non-significant **[C]**. **[E] Fasting plasma insulin levels.** A significant reduction in plasma insulin was observed in all the treated groups compared to HFD-fed mice. **[F] Fasting plasma leptin levels.** A significant reduction in plasma leptin was observed in M & M+G treated groups compared to HFD-fed mice. **[G] Fasting plasma melatonin levels.** A significant increase in plasma melatonin was observed in M & M+G treated groups compared to HFD-fed mice. (\**p*<0.05, \*\**p*<0.01, \*\*\**p*<0.001 vs. NCD; ^@^*p*<0.05, ^@@^*p*<0.01, ^@@@^*p*<0.001 vs. HFD; ^&^*p*<0.05, ^&&&^*p*<0.001 vs. Melatonin; ^#^*p*<0.05, ^##^*p*<0.01, ^###^*p*<0.001 vs. GABA) (n=5-6/group).

#### 3.2.3 Assessment of key glucoregulatory enzyme gene expression & specific activity and liver glycogen content

GCK enzyme activity was significantly increased in the M+G group (*p*<0.05) compared to the HFD group indicating increased glucose uptake. *G6Pase* expression was significantly reduced in all the treated groups (M, G, *p*<0.05; M+G, *p*<0.01) compared to the HFD group, indicating increased glucose uptake, and *GLUT2* expression was significantly reduced in GABA and M+G groups (G, M+G, *p*<0.05) as compared to HFD group. Furthermore, the gene expression and activity of FBPase and PEPCK were significantly increased in the HFD group (*p*<0.05 and *p*<0.01, respectively) compared to NCD groups and significantly reduced in M & M+G and G & M+G groups, respectively, suggesting reduced gluconeogenesis. Additionally, there was a significant reduction in the gene expression (M, *p*<0.05; G, *p*<0.05; M+G, *p*<0.05) and activity of GP (M, *p*<0.001; G, *p*<0.01; M+G, *p*<0.001) in all the drug-treated groups as compared to HFD indicating reduced glycogenolysis. No significant change in *GS* expression was observed in any drug-treated group (*p*>0.05) compared to HFD. However, glycogen content in all the drug-treated groups (S, M, S+M, *p*<0.001) indicated an increase in glycogenesis (Figure 5A-L).

**Figure 5:**
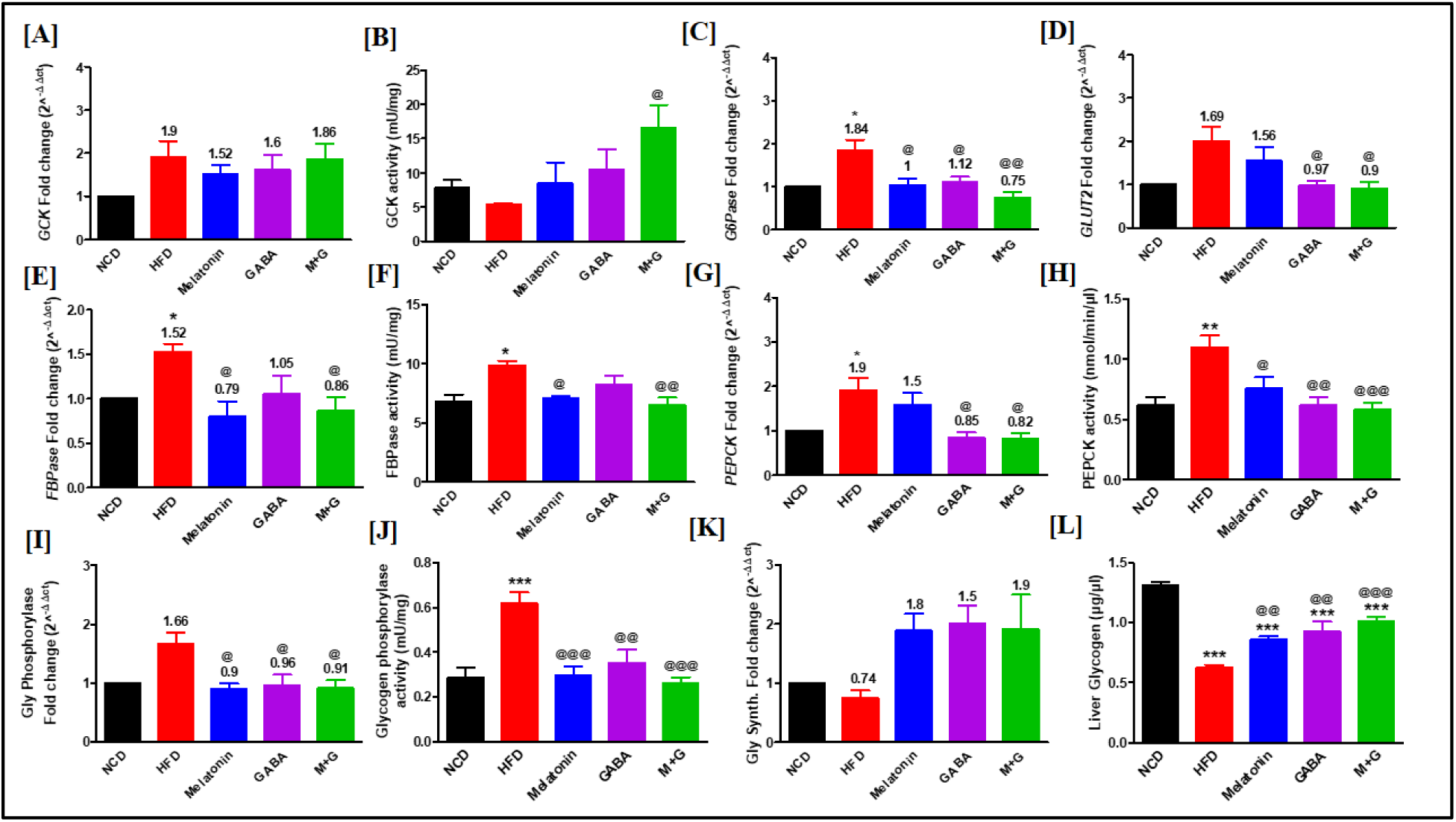
Gene expression and enzyme activities of glucoregulatory enzymes and glycogen content in the liver. **[A-B] *GCK* mRNA fold-change and activity.** No significant change in *GCK* expression was observed in any of the treated groups compared to HFD-fed mice. Intriguingly, its activity was significantly increased in the M+G treated group. **[C] *G6Pase* mRNA fold change.** *G6Pase* expression significantly increased in HFD, decreasing significantly in all the drug-treated groups. **[D] *GLUT2* mRNA fold change.** A significant decrease in *GLUT2* expression was observed in the G and M+G treated groups compared to HFD: [E-F] *FBPase* mRNA fold-change and its activity. *FBPase* gene expression and its activity were significantly increased in HFD, whereas in the M and M+G treated groups, it was reduced significantly—**[G-H] *PEPCK* mRNA fold-change and activity.** *PEPCK* expression and activity were significantly increased in HFD. In contrast, its fold change was significantly decreased in the G and M+G treated groups, and activity was decreased in all the drug-treated groups. **[I-J] *GP* mRNA fold-change and activity.** All the drug-treated groups observed a significant decrease in GP expression and activity. **[K] *GS* mRNA fold change**. No significant change in *GS* expression was observed in any of the treated groups compared to HFD-fed mice. **[L] Liver glycogen content (GC).** GC was significantly reduced in HFD, whereas it was increased significantly in all the drug-treated groups. (\**p*<0.05, \*\**p*<0.01, \*\*\**p*<0.001 vs. NCD; ^@^*p*<0.05, ^@@^*p*<0.01, ^@@@^*p*<0.001 vs. HFD) (n=5-6/group).

#### 3.2.4 Assessment of mitochondrial biogenesis markers in skeletal muscle, lipid metabolism markers, and absolute gene quantification of *MTNR1B* and *GLUT4* in adipose tissue

Expression of crucial markers for mitochondrial biogenesis, i.e. *Sirtuin 1* (*SIRT1*) and *Peroxisome proliferator-activated receptor gamma coactivator 1-alpha* (*PGC1α*), showed a significant increase in all the drug-treated groups as compared to HFD (SIRT1: G, *p*<0.01; M, M+G, *p*<0.001; PGC1α: M, G, *p*<0.05; M+G, *p*<0.01). Lipid metabolism markers, i.e. *Acetyl-CoA carboxylase* 1 (*ACC1*) expression was reduced in both HFD and the drug-treated groups (M, G, M+G, *p*<0.001), whereas *Adipose triglyceride lipase* (*ATGL*) expression was significantly reduced in all the drug-treated groups (G, *p*<0.05; M, M+G, *p*<0.001) as compared to HFD, suggesting improved lipid metabolism (Figure 6A-D). Additionally, we studied the absolute gene expression of *MTNR1B* and *GLUT4* in adipose tissue by ddPCR. The results show an increased copy number/µl of the mentioned genes in the HFD group compared to the NCD group, suggesting insulin resistance. In contrast, the copy number/µl of *MTNR1B* and *GLUT4* were reduced in all the treated groups, suggesting reduced insulin resistance (Figure 6E-F).

**Figure 6:**
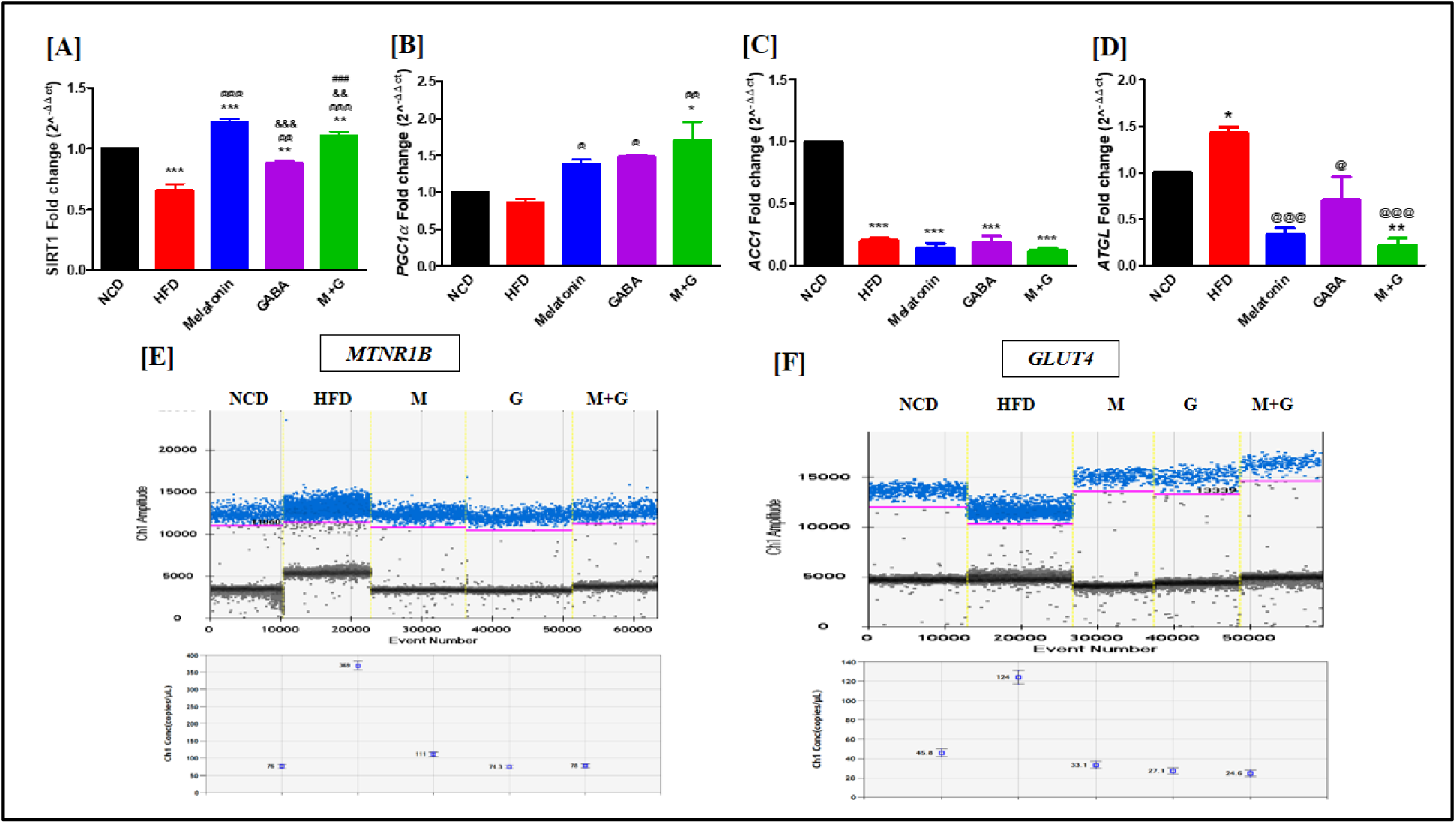
Gene expression of mitochondrial biogenesis markers in skeletal muscle, lipid metabolism markers, and absolute gene quantification of *MTNR1B* and *GLUT4* in adipose tissue. [A-B] mRNA fold change of *SIRT1* and *PGC1a* in skeletal muscle. A significant reduction in *SIRT1* mRNA levels was observed in HFD, whereas all the drug-treated groups showed a significant increase in SIRT1 and PGC1a levels—**[C-D] mRNA fold change of *ACC1* and *ATGL* in adipose tissue.** There was no change in *ACC1* expression, whereas a significant reduction in *ATGL* gene expression was observed in all the drug-treated groups compared to HFD—**[E-F] absolute gene quantification of *MTNR1B* and *GLUT4* in adipose tissue**. Gene expression of *MTNR1B* and *GLUT4* was higher in the HFD group than in the NCD group. In contrast, it was reduced in all the drug-treated groups compared to the HFD groups, as represented by a change in the amplitude of positive droplets (blue) and the number of copies/μl. Blue droplets: positive for the target gene; Grey droplets: negative for the target gene; Pink line: threshold intensity to discriminate positive and negative droplets (n=3/group). (\**p*<0.05, \*\**p*<0.01, \*\*\**p*<0.001 vs. NCD; ^@^*p*<0.05, ^@@^*p*<0.01, ^@@@^*p*<0.001 vs. HFD; ^&&^*p*<0.01, ^&&&^*p*<0.001 vs. Melatonin; ^###^*p*<0.001 vs. GABA) (n=6-8/group).

#### 3.2.5 Estimation of Mitochondrial Respiratory Control Ratio (RCR) of Complexes I-IV in Skeletal Muscle

The RCR of state 3/state 4 for mitochondrial complexes I, II, III, and IV (*p*<0.001, *p*<0.001, *p*<0.01, *p*<0.05, respectively) was significantly reduced in the HFD group as compared to NCD. A significant increase in the RCR was observed in all the treated groups for CIII (M, *p*<0.05; G, *p*<0.05; M+G, *p*<0.01) and CIV (M, *p*<0.01; G, *p*<0.05; M+G, *p*<0.01). However, the RCR of state 3/state 4 for CI was significantly increased only in the M+G (*p*<0.05) and for CII in the M (*p*<0.05) and M+G (*p*<0.001) treated group (Figure 7A-D).

**Figure 7:**
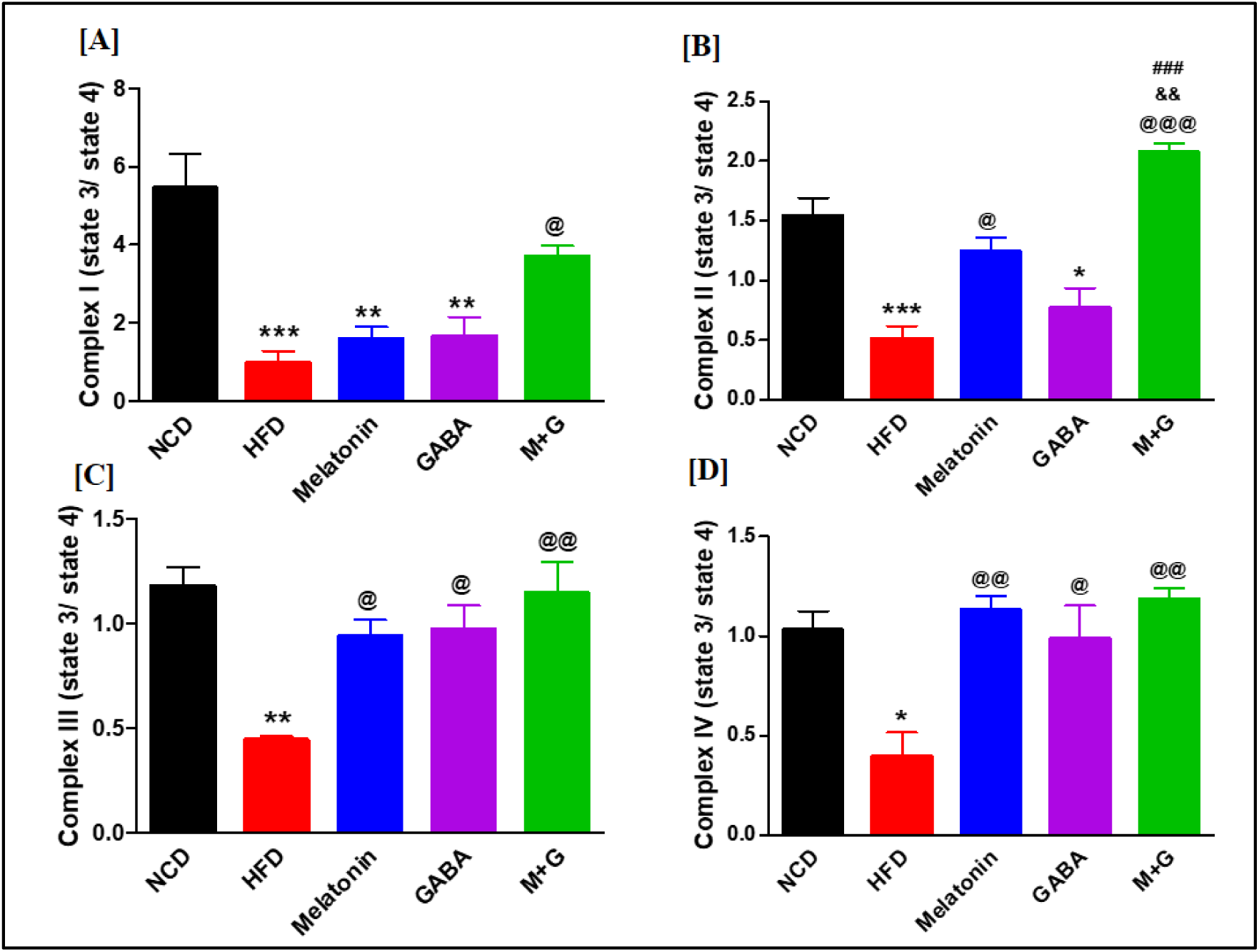
Mitochondrial respiratory control ratio (state 3/state 4) in Skeletal muscle. **[A] Complex I.** RCR was significantly reduced in the HFD group as compared to NCD and was significantly increased in the M+G treated group **[B] Complex II.** RCR was reduced significantly in the HFD group compared to NCD and increased significantly in the M and M+G treated groups. **[C-D] Complex III-IV.** RCR was significantly reduced in the HFD group compared to NCD and increased significantly in all the drug-treated groups. (\**p*<0.05, \*\**p*<0.01, \*\*\**p*<0.001 vs. NCD; ^@^*p*<0.05, ^@@^*p*<0.01, ^@@@^*p*<0.001 vs. HFD; ^&&^*p*<0.01 vs. Melatonin; ^###^*p*<0.001 vs. GABA) (n=3/group).

#### 3.2.6 Protein Expression Analysis for Insulin Signaling Pathway in Skeletal Muscle

Western blot analysis of key proteins involved in the insulin signalling pathway showed that IR1β (*p*<0.05), pAkt Ser473/Akt (*p*<0.05), and GLUT4 (*p*<0.05) levels were significantly downregulated in HFD-fed mice, while pIRS Ser307/IRS (*p*<0.05) levels significantly upregulated in HFD (*p*<0.05) as compared to NCD, which is suggestive of peripheral insulin resistance. In the drug-treated groups, there was an upregulation of IR1β (M, *p*<0.05; G, *p*<0.05; M+G, *p*<0.01), and pAkt/Akt levels (M, *p*<0.05; M+G, *p*<0.01), as compared to HFD group. Moreover, the levels of pIRS/IRS were significantly downregulated in M and M+G groups (M, *p*<0.05; M+G, *p*<0.05), suggesting improved insulin sensitivity. Intriguingly, GLUT4 levels were significantly upregulated only in M+G treated groups (*p*<0.05) as compared to the HFD group (Figure 8A-E).

**Figure 8:**
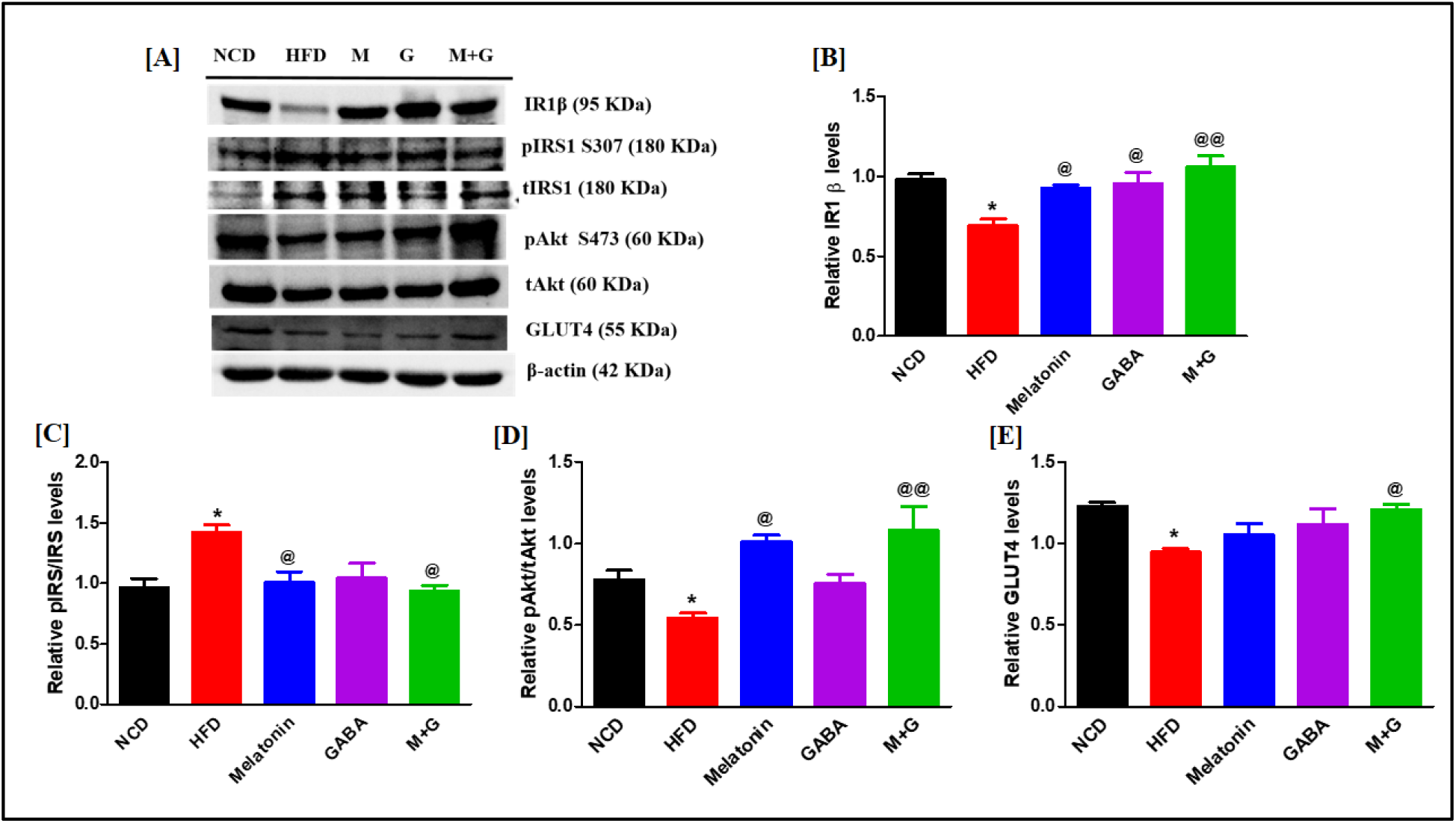
Protein expression of the insulin signalling pathway in skeletal muscle. **[A]** Representative western blot images. **[B] Insulin Receptor 1β.** IR1β protein expression was significantly reduced in HFD compared to NCD, whereas all the drug-treated groups observed a significant increase in IR1β. **[C] pIRS/IRS.** pIRS/IRS levels were significantly increased in HFD and were reduced significantly in M and M+G treated groups. **[D] pAkt/Akt.** pAkt/tAkt levels significantly decreased in HFD while increasing significantly in the M and M+G treated groups. **[E] GLUT4**. GLUT4 protein expression was significantly decreased in the HFD while increasing significantly in the M+G treated group. (\**p*<0.05 vs. NCD; ^@^*p*<0.05, ^@@^*p*<0.01 vs. HFD) (n=4/group).

#### 3.2.7 Assessment of β-cell Mass and Islet Number in Pancreas

IHC analysis of the pancreatic tissue showed that β-cell mass was significantly increased in the HFD group compared to NCD (*p*<0.001). All the drug-treated groups (G, *p*<0.01; M, M+G, *p*<0.001) showed a significant decrease in β-cell mass compared to the HFD group. Intriguingly, the islet number per pancreatic section was also significantly increased in the HFD group (*p*<0.001) as compared to NCD; however, it was notably reduced in all the drug-treated groups (M, G, M+G, *p*<0.001) as compared to HFD group, revealing a reduction in beta-cell hyperplasia and hypertrophic condition as shown in Figure 9A-C.

**Figure 9:**
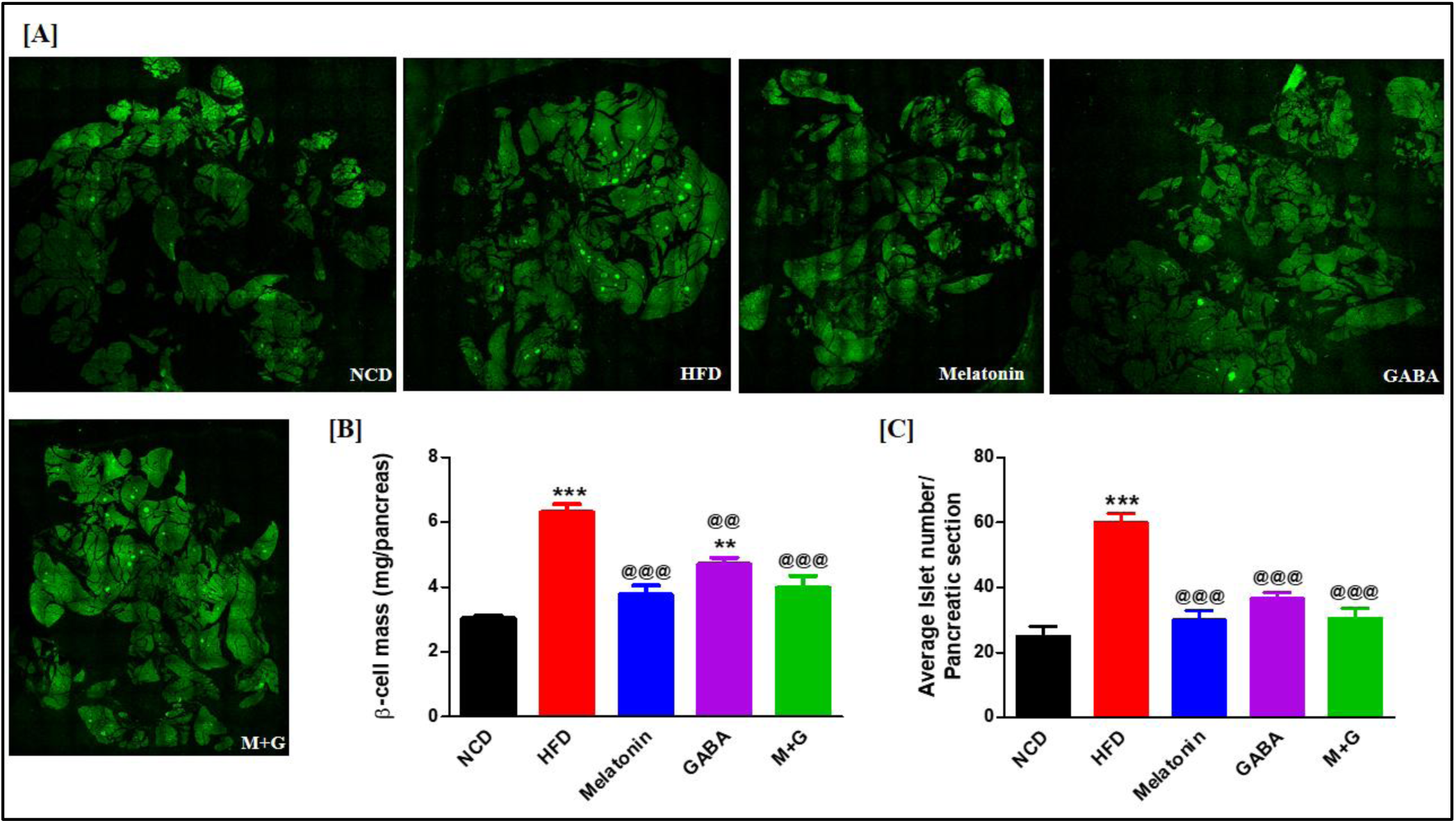
Pancreatic β-cell mass and Islet number. **[A]** Immunofluorescence staining for insulin (green) in mouse pancreatic tissue. **[B] Pancreatic β-cell mass**: β-cell mass was significantly increased in the HFD group while it was reduced significantly in all the treated groups compared to the HFD group. **[C] Islets Number per pancreatic section.** Islet number was significantly increased in the HFD group compared to the NCD group, while it was reduced significantly in all the drug-treated groups compared to the HFD group. Magnification-10X. (\*\**p*<0.01, \*\*\**p*<0.001 vs. NCD; ^@@^*p*<0.01, ^@@@^*p*<0.001 vs. HFD) (n=3/group).

## 4. Discussion

The management of diabetes mellitus involves two main approaches: i) increasing the survival rate and proliferation of endogenous pancreatic β-cells with immunosuppressive effects and ii) increasing insulin-mediated glucose uptake by the peripheral tissues. However, none of the current therapies effectively achieve both goals [30]. Recent studies suggest that melatonin and GABA both individually can stimulate β-cell replication, protect β-cells against apoptosis, attenuate insulitis, regulate the islet-cell function and glucose homeostasis, improve peripheral insulin sensitivity and suppress detrimental immune reactions [4,5,21–24]. Hence, the present study investigated the potential beneficial effects of melatonin administration combined with GABA in T1D and T2D rodent models. Our results show that monotherapies and combination therapy can sub-additively ameliorate T1D and T2D manifestations. In the T1D model, treatments could induce β-cell regeneration (BrdU^+^, ARX^+^, PAX4^+^ β-cells), reduce β-cell apoptosis (TUNEL+), increase insulin levels, lower FBG, and improve impaired glucose tolerance. In addition, it improves glucose and lipid metabolism by ameliorating hyperglycemia, hyperinsulinemia, hyperleptinemia, and dyslipidemia. Moreover, it preserves β-cell mass and islet number and increases mitochondrial biogenesis and insulin sensitivity in skeletal muscle and adipose tissue in the T2D model.

Due to the lack of NGN3 expression in all the drug-treated groups, we suggest that β-cell neogenesis was absent in all the groups. Further, PAX4^+^ ARX^+^ insulin^+^ co-positive cells were observed in all drug-treated groups, indicating the presence of α-to β-cell transdifferentiation. Moreover, PAX4 also defines a proliferating β-cell population [31,32], which was increased in all drug-treated groups. However, lineage-tracing experiments are required to substantiate our results on β-cell transdifferentiation. Furthermore, our results suggest that β-cell proliferation (INS^+^ BrdU^+^ co-positive cells) was significantly increased, and apoptosis (INS^+^ TUNEL^+^ co-positive cells) was significantly reduced in all the treated groups. Interestingly, our study also reveals increased glycolysis, glycogen synthesis, and glycogen content and reduced gluconeogenesis, glycogenolysis, and *GLUT2* expression in the liver, suggesting improved glucose metabolism in all the drug-treated groups. We have observed decreased *ATGL* (lipolytic) transcript levels in adipose tissue with no change in *ACC1* (lipogenic) in all the drug-treated groups. In addition, increased *SIRT1* and *PGC1-α* (mitochondrial biogenesis genes) transcript levels in the skeletal muscle indicate improved lipid metabolism and insulin sensitivity in the respective tissues. Our mitochondrial studies suggest increased respiratory chain complexes I-IV activities in skeletal muscle in all the drug-treated groups, presumably by improving *SIRT1* and *PGC1-α* levels upon drug treatment. Atkinson *et al.* reported that overexpression of *GLUT4* increases insulin sensitivity and glucose tolerance in HFD-fed mice [33], and its expression changes with short-term and long-term HFD feeding [34]. In support of these studies, our ddPCR results revealed an increased expression of *MTNR1B* and *GLUT4* in the HFD group, indicating insulin resistance.

In contrast, the levels in the drug-treated groups in adipose tissue were restored. It is possible that the overexpression was to compensate for insulin resistance in the HFD group, which was then restored in the treatment groups showing increased insulin sensitivity. In addition, *MTNR1B* overexpression in adipose tissue leads to increased lipid levels and adipocytes [35], which is supported by our results showing increased TG levels and fat in the HFD group. At the same time, we observe restored expression of *MTNR1B* expression and lipid levels in the drug-treated groups. Overnutrition causes mTOR/S6K1 signalling activation and impairs insulin signalling by increased serine phosphorylation of IRS1 [36,37]. We report a significant decrease in serine phosphorylation of IRS1 and an increase in IR1β, pAKT, and GLUT4 expression in all drug-treated groups, suggesting an improving insulin signalling pathway in skeletal muscle.

Besides the pineal gland, melatonin is locally synthesized in tissues like the retina, pancreas, adipose tissue, and liver. It mediates its action via melatonin receptor 1A (MT1) and melatonin receptor 1B (MT2), regulating metabolic functions. By binding to Gq-coupled MT2 receptors, melatonin stimulates the activity of PLC and IP3, leading to insulin secretion [38]. Chan *et al.* show that melatonin treatment increases insulin secretion and promotes β-cell survival via decreased c-JUN N-terminal kinase (JNK) activation in human pancreatic islets [39]. Melatonin inhibits lipolysis and fatty acid transport in rat adipocytes by inhibiting the cAMP–PKA pathway [40]. In hepG2 cells, melatonin stimulates glycogen synthesis via protein kinase C, zeta (PKCζ), Akt, and glycogen synthase kinase-3β (GSK3β) [41], besides reducing gluconeogenesis in rats [42]. The findings of melatonin regulating glucose metabolism via increased PI3K-Akt activity [35,43] support our findings. Further, it is reported that melatonin activates IRS1–PI3K–PKCζ pathway and stimulates glucose uptake in mouse skeletal muscle cells [44], while it is reduced in MT1 knockout mice [45].

GABA has a proven role in islet-cell hormone homeostasis, preserving the β-cell mass, suppressing immune reactions and consequent apoptosis [46]. The GABAergic system functions in many peripheral tissues like the pancreas and liver. GABA_A_R regulates PI3K/Akt activities in the liver that maintain hepatocyte survival and PGC-1α expression. Thus, GABA is one of the factors responsible for regulating hepatic glucose metabolism, i.e., gluconeogenic and glycogenolytic pathways [47]. Fibroblast growth factor 21 (FGF-21), predominantly expressed in the liver under normal conditions, induces PGC-1α expression, leading to the regulation of lipolysis in the adipose tissue, forming liver-adipose tissue crosstalk [48]. Choi *et al.* have reported that GABA-enriched fermented sea tangle promotes brain-derived neurotrophic factor-mediated muscle growth and lipolysis in middle-aged women [49]. Previously, it was reported that T2D subjects had impaired mitochondrial TCA cycle flux in skeletal muscle [50,51]. In a series of enzymatic reactions, the TCA cycle generates the reducing equivalents NADH and FADH2 required to transfer electrons to the mitochondrial respiratory chain [52]. It could be one possible reason for impaired mitochondrial respiration in T2D conditions. Increased PGC1α activates branched-chain amino acid (BCAAa) metabolism, fatty acid oxidation, and the TCA cycle in the skeletal muscle [53–55]. We found that the GABA-treated group increased *SIRT1* and *PGC1α* expression. Thus, we hypothesized that GABA could improve mitochondrial biogenesis and oxygen consumption rate, possibly by regulating *SIRT1* and *PGC1α*. GABA via GABA_A_R and GABA_B_R promotes the phosphorylation of CREB (cAMP response element-binding protein) [21], a key transcription factor responsible for the maintenance of insulin gene transcription and β-cell survival in rodent and human islets [56]. Among targets of CREB, the insulin receptor substrate-2 gene (IRS-2) is crucial in regulating β-cell mass and function [21]. GABA_A_R regulates PI3K/Akt activities in the pancreas that induce insulin secretion, beta-cell survival, and regeneration [57]. Possibly, GABA has an anti-apoptosis effect due to the PI3K/Akt pathway, as this pathway can regulate both anti- and pro-apoptotic pathways. Lin *et al.* have reported that melatonin suppresses autoimmune recurrence in NOD mice by inhibiting the proliferation of Th1 cells and decreasing the production of pro-inflammatory cytokines [58]. GABA regulates cytokine secretion from human PBMCs, suppresses β-cell-reactive CD8^+^ CTLs, and increases Regulatory T-Cell in T1D models [57]. GABA’s anti-inflammatory and immunomodulatory actions are governed by SIRT1-mediated suppression of the NF-κB pathway, which further protects pancreatic β-cells against apoptosis [59].

These effects of melatonin and GABA were first observed in mice [23,60,61], and it also appears valid in humans as demonstrated in vitro and xenotransplanted human islets [4,21,62]. We assume that combination therapy sub-additively ameliorates T1D and T2D diabetic manifestations, as monotherapies are as effective as combination therapy. Toews and Bylund reported that if the sum of the responses to the two drugs individually is greater than the maximal response possible by the system, then subadditivity occurs without any true drug interaction [63]. Further, we assume that the reasons for getting a sub-additive effect could be i) the selected dose of M and G might be the maximum effective dose, and ii) there is no true interaction between M and G in the system. The limitation of our study is that we have not looked into the mechanistic aspect of the M and G mode of action. Unlike G, most of the pathways by which melatonin exerts its pleiotropic effects are well known [16,64]. However, we summarized the known and possible hypothesized modes of action of M and G based on the studies that support our findings. The possible mode of action of M and G and their combinatorial effects on the amelioration of diabetes manifestations in peripheral tissues (pancreas, liver, skeletal muscle, and adipose tissue) are shown in Fig. S3.

## 5. Conclusion

Our studies suggest that melatonin and GABA sub-additively ameliorate T1D and T2D in mouse models by inducing β-cell regeneration and survival, improving glucose and lipid metabolism, and increasing insulin sensitivity in peripheral tissues. Overall, this knowledge can eventually be used in future translational research, leading to the development of targeted drug therapy for diabetes. It is also essential to consider the dosage of both the drug, melatonin treatment (acute/chronic) and the relative timing of its intake in relation to glycemic challenges in humans.

## Statements and Declarations Ethical Statement

The submitted manuscript is not being considered for publication in any other journal, and the authors declare that there are no financial/competing interests. All authors have read and approved all versions of the manuscript, its content, and its submission to the *Journal of Endocrinological Investigation*.

## Ethical approval

All the experimental procedures were conducted as per the Purpose of Control and Supervision of Experiments on Animals (CPCSEA) guidelines and were approved by the Institutional Animal Ethical Committee (IAEC) (MSU/BC/02/2019; MSU/BC/09/2019).

## Data Availability

The datasets generated during and/or analyzed during the current study are included in this article and are not publicly available due to privacy policy. However, they are available from the corresponding author upon reasonable request.

## Authors’ Contributions

RB conceived the idea. NP designed and performed the experiments, data acquisition, and data analysis and wrote the manuscript. RP and NR helped acquire data, review, and edit the manuscript. RB, MD and AVR contributed to the critical revision and approval of the manuscript.

## Acknowledgements

We thank Dr. Murali Chilkapati and Mrs. Vaishali Kailaje, the digital imaging facility at The Advanced Centre for Treatment, Research and Education in Cancer (ACTREC), Mumbai, India, for providing the conference facilities. We thank Mr. Darshan Mehta for helping us use the conference facility at ACTREC. We thank Dr. Gurprit Bhardwaj and Dr. Dhruv Gohel, Department of Biochemistry, MSU Baroda, India, for helping us in sample preparation for western blot analysis and ddPCR study, respectively. We also thank Dr. Sayantani Pramanik Palit, Ms. Naisargi Patel, and Ms. Satyashree Shetty for their help. NP thanks the Gujarat government for awarding the SHODH fellowship. RP thanks CSIR, New Delhi, India, for awarding SRF. NR thanks UGC-NFST, New Delhi, India, for awarding JRF and SRF.

