## Supplementary Figures & Table for "Melatonin and GABA monotherapies are as efficacious as combination therapy in managing T1D and T2D: *In-vivo* studies on experimental diabetic models"

**Supporting Information:**

**Figure S1. The experimental timeline and over strategy for T1D model generation.**

**~**7-8 weeks old Male BALB/c mice were taken and given one-week acclimatization and then given five consecutive intraperitoneal injections of 50 mg/kg body weight STZ. Control mice were given sodium citrate buffer as vehicle control for STZ. BW and FBG levels were monitored twice week. Once T1D was confirmed, these animals were divided randomly into four groups for treatment for six weeks. At the end of treatment, IPGTT and IPITT were carried. Afterward, the animals were sacrificed, and tissues were harvested for further analysis.

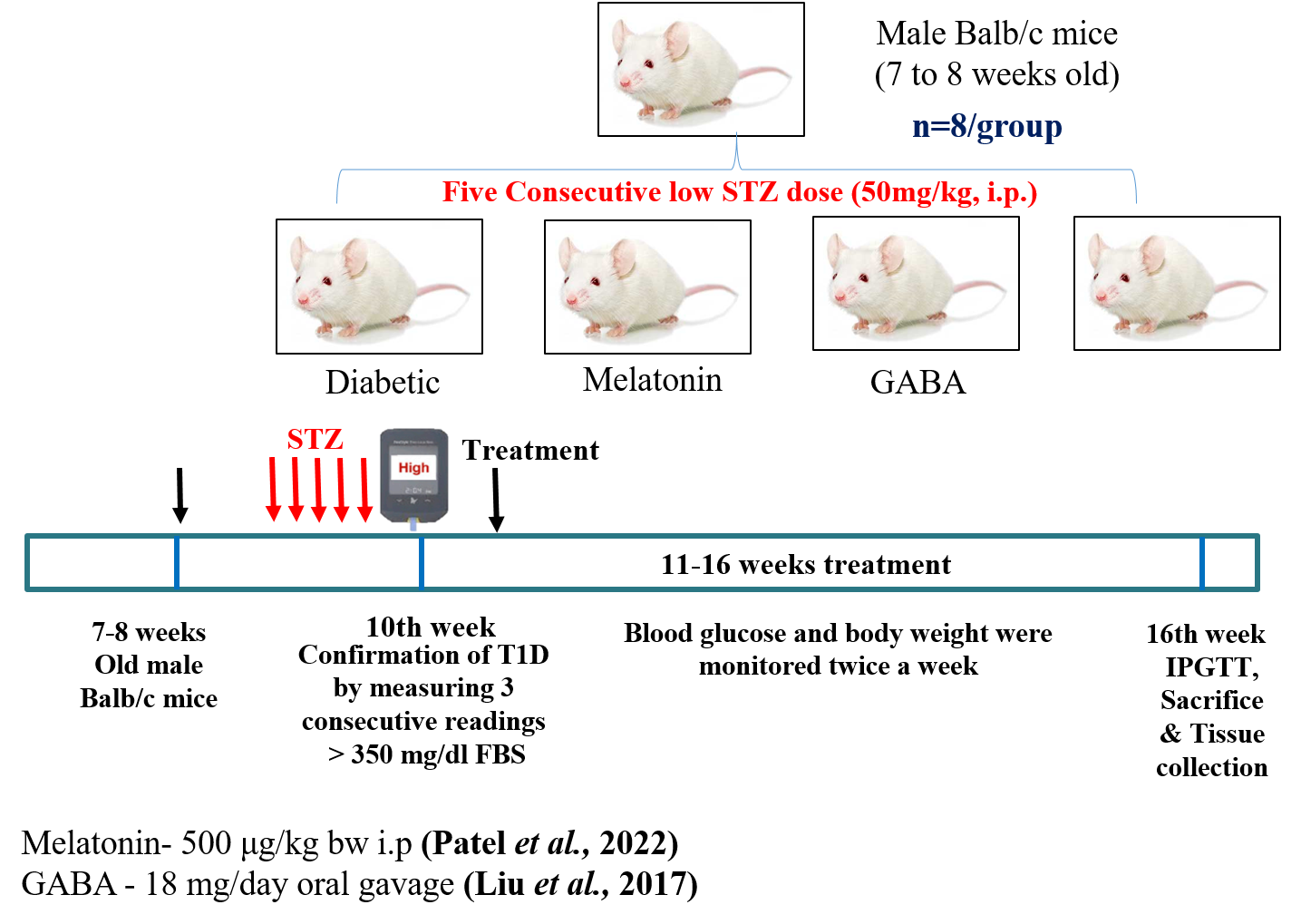

**Figure S2. The experimental timeline and over strategy for T2D model generation.**

**~**6-7 weeks old Male C57BL/6 mice were procured and given one-week acclimatization and then proceeded for 30 weeks of HFD to induce obesity and insulin resistance. A non-diabetic control group was fed with chow diet. The food intake, water consumption, BW, and FBG levels were monitored weekly. Once obesity-induced T2D was confirmed, these animals were divided randomly into four groups for treatment for six weeks. At the end of treatment, IPGTT and IPITT were carried. Afterward, the animals were sacrificed, and tissues were harvested for further analysis.

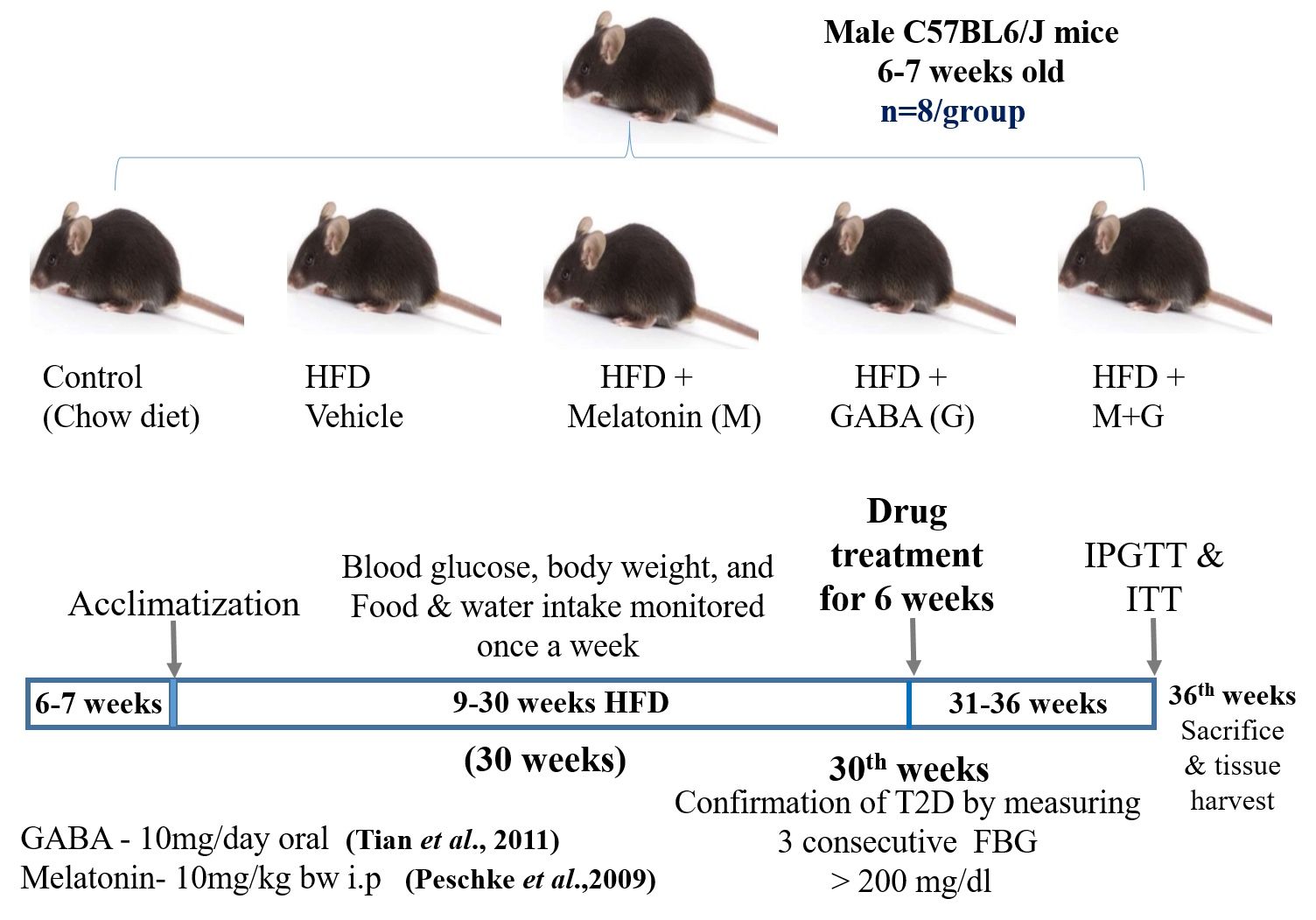

**Figure S3. Possible mode of action of Melatonin and GABA and their combination in the amelioration of diabetes manifestations in peripheral tissues.**

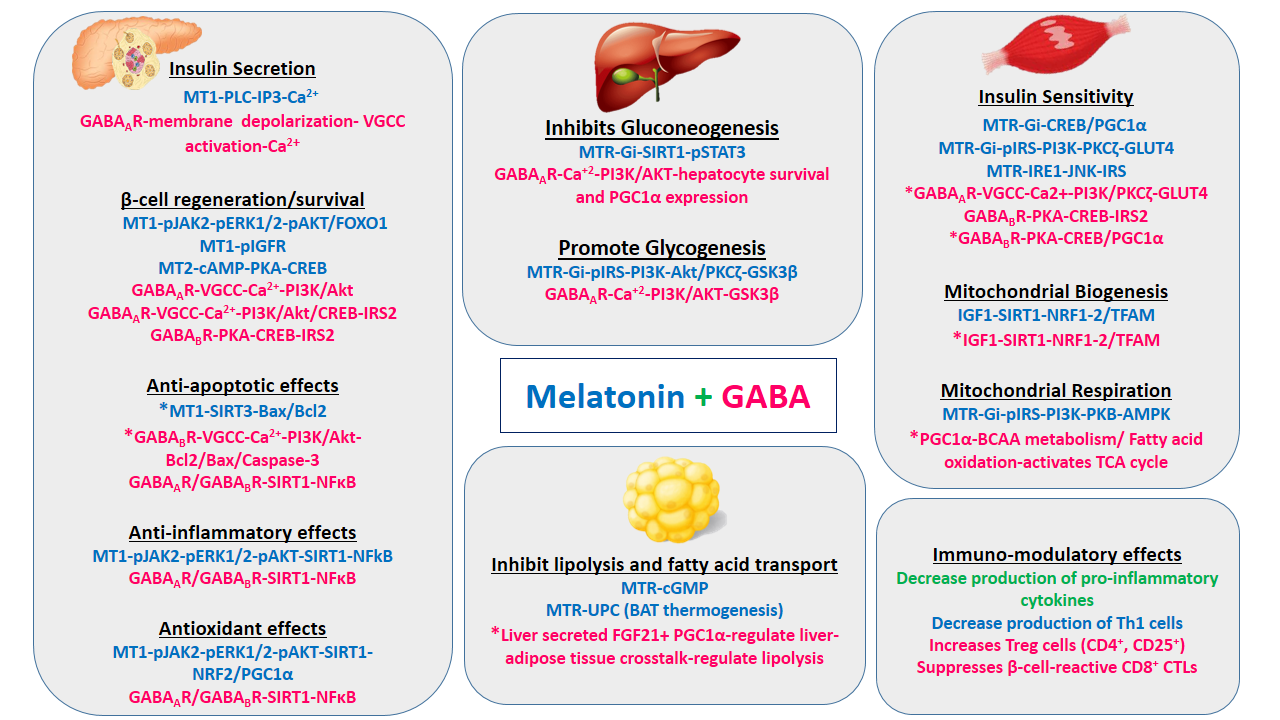

Melatonin mediates its action [16,64] on insulin secretion (MT1-IP3-Ca2+), β-cell regeneration and survival (cAMP-PKA-CREB; pJAK2-pERK1/2-pAKT/FOXO1), insulin sensitivity (MTR-Gi-CREB/PGC1α; pIRS1-GLUT4; IRE1-JNK-IRS1), glycogenesis (MTR-Gi-pIRS1-GSK3β), inhibiting gluconeogenesis (MTR-Gi-SIRT1-pSTAT3), mitochondrial biogenesis (IGF1-SIRT1-NRF1/2-TFAM), mitochondrial respiration (MTR-Gi-pIRS-PI3K-PKB-AMPK), inhibiting lipolysis (MT-cGMP), BAT thermogenesis (MTR-UCP). It also mediates antioxidant effects (MT1-pAKT-NRF2/PGC1α), anti-apoptotic effects (*MT1-SIRT1-BAX/BCL2), and anti-inflammatory effects by reducing Th1 cells and MT1-pJAK2-pERK1/2-pAKT-SIRT1-NFκB pathway. Melatonin decreases the production of Th1 cells. GABA mediates its action [21,47,48,51,53-55,59] on insulin secretion (GABA_A_R -VGCC activation-Ca^2+^), insulin sensitivity (GABA_B_R-PKA-CREB-IRS2; *GABA_B_R-PKA-CREB/PGC1α; *GABA_A_R-VGCC-Ca2+-PI3K/PKCζ-GLUT4) β-cell regeneration and survival (GABA_A_R-VGCC activation-Ca^2+^-pI3K/Akt; VGCC-Ca^2+^-pI3K/Akt /CREB-IRS2; GABA_B_R-PKA-CREB-IRS2), anti-apoptotic effects (GABA_A_R/GABA_B_R-SIRT1-NFκB; *GABA_B_R-VGCC-Ca^2+^-pI3K/Akt-Bcl2/Bax/Caspase-3), anti-inflammatory effects (GABA_A_R/GABA_B_R-SIRT1-NFκB), glycogenesis (GABA_A_R-Ca^+2^-PI3K/AKT-GSK3β), inhibiting gluconeogenesis (*GABA_A_R-Ca^+2^-PI3K/AKT-hepatocyte survival and PGC1α), inhibiting lipolysis (Liver secreted FGF21+ PGC1α-regulate liver-adipose tissue crosstalk-regulate lipolysis), mitochondrial biogenesis (*IGF1-SIRT1-NRF1-2/TFAM), mitochondrial respiration (*PGC1α-BCAA metabolism/ Fatty acid oxidation-activates TCA cycle). GABA suppresses β-cell-reactive CD8+ CTLs and increases Treg cells (CD4+, CD25+). Both melatonin and GABA have anti-inflammatory effects by reducing the production of pro-inflammatory cytokines.

AMPK: AMP-activated protein kinase; GABA_A_R: GABA receptor  type A; GABA_B_R: GABA receptor  type B; MTR: Melatonin receptor; MT1: Melatonin receptor 1; MT2: Melatonin receptor 2; BAX: BCL2 associated X protein; BCL2: B-cell lymphoma 2; cGMP: Cyclic guanosine monophosphate; Caspase-3: cysteine–aspartic acid protease; CREB: -response element binding protein; FOXO1: Forkhead box protein O1; GLUT4: Glucose transporter type 4; IGF1: Insulin-like growth factor 1; IP3: Inositol trisphosphate; IRE1: Inositol-requiring enzyme 1; IRS2: Insulin receptor substrate 2; NFκB: Nuclear Factor kappa-light-chain-enhancer of activated B cells; NRF1/2: Nuclear respiratory factor 1 and 2; PDX1: Pancreatic and duodenal homeobox 1; PGC1α: Peroxisome proliferator-activated receptor gamma coactivator 1-alpha; PI3K: Phosphoinositide 3-kinase; PKA: Protein kinase A; PKB/AKT: Protein kinase B; PKCζ: Protein kinase C zeta; PLC: Phospholipase C; pERK1/2: Phospho-Extracellular signal-regulated protein kinases 1 and 2; pIGFR: Phospho-Insulin-like growth factor 1 receptor; pIRS1: Phospho-Insulin receptor substrate 1; pJAK2: Phospho-Janus kinase 2; pSTAT3: Phospho-Signal transducer and activator of transcription 3; SIRT1/3: Sirtuin 1/3; GSK3B: Glycogen Synthase Kinase 3 Beta; JNK: c-Jun N-terminal kinase; TFAM: transcription factor A, mitochondrial; BCAA: Branched-chain amino acids; VGCC: Voltage-gated calcium channels; TCA: The tricarboxylic acid; Th1: T helper type 1; Tregs: Regulatory T cells; CTLs: cytotoxic T cell. *Hypothesized pathway.

**Table S1.** Primers used for the transcript analysis.

| **Gene**  **Primer** | **Sequence (5'-3')** | **Annealing**  **Temperature** | **Amplicon**  **Size (bp)** | **Tissue** |
| --- | --- | --- | --- | --- |
| *Glucokinase*  (*GCK*) | **FP:** AGGAGGCCAGTGTAAAGATGT  **RP:** TCCCAGGTCTAAGGAGAGAAA | 56°C | 90bp | Liver |
| *Phosphoenol-*  *pyruvate carboxykinase* (*PEPCK*) | **FP:** CTGCATAACGGTCTGGACTTC  **RP:** CAGCAACTGCCCGTACTCC | 65°C | 151bp |  |
| *Fructose bis-*  *phosphatase 1*  *(FBP1)* | **FP:** GCATCGCACAGCTCTATGGT  **RP:** CTCAGGTTCGATTATGATGGC | 59°C | 170 bp |  |
| *Glucose-6-*  *phosphatase*  (*G6Pase*) | **FP:** CTGTTTGGACAACGCCCGTAT  **RP:** AGGTGACAGGGAACTGCTTTA | 56°C | 91bp |  |
| *Glucose transporter 2* (*GLUT2)* | **FP:** CTTGGAAGGATCAAAGCAATG  **RP:** CAGTCCTGAAATTAGCCCAC | 60°C | 150bp |  |
| *Glycogen Synthase* (*GS*) | **FP:** ACCAAGGCCAAAACGACAG  **RP:** GGGCTCACATTGTTCTACTTG | 61°C | 102bp |  |
| *Glycogen Phosphorylase*  (*GP*) | **FP:** GAGAAGCGACGGCAGATCA  **RP:** CTTGACCAGAGTGAAGTGCA | 65°C | 102bp |  |
| *Sirtuin 1*  (*SIRT-1*) | **FP:** GATGAAGTTGACCTCCTCA  **RP:** GGGTATAGAACTTGGAATTAG | 61°C | 86bp | Skeletal muscles  (SK) |
| *Peroxisome proliferator-activated receptor gamma coactivator 1-alpha*  (*PGC-1α*) | **FP:** AGCCGTGACCACTGACAACGA  **RP:** GTAGCTGAGCTGAGTGTTGGC | 69°C | 129bp |  |
| *Adipose triglyceride lipase*  (*ATGL*) | **FP:** CAACGCCACTCACATCTACGG  **RP:** GGACACCTCAATAATGTTGGCAC | 61.8°C | 158bp | Adipose  Tissue  (AT) |
| *Acetyl-CoA*  *carboxylase* 1 (*ACC-1*) | **FP:** ACGCTCAGGTCACCAAAAAGAAT  **RP:** GTAGGGTCCCGGCCACAT | 57°C | 70bp |  |
| *Melatonin receptor B*  (*MTNR1B*) | **FP:**  TTGTGATGGGCCTGAGTGTC  **RP:**  AGCCAGACGAGGCTGATGTA | 60°C | 145bp |  |
| *Glucose transporter 4* (*GLUT4*) | **FP:**  TAGGAGCTGAGGGTTGGCTA  **RP:** TGCTCCAGTAGGCCGTAAAC | 60°C | 111bp |  |
| *GAPDH* | **FP:** AGGTCGGTGTGAACGGATTTG  **RP:** TGTAGACCATGTAGTTGAGGT | 56°C | 123bp | Liver, SK, AT |

FP: Forward Primer; RP: Reverse Primer; bp: base pair

**Table S2.** Antibodies used for the immunoblot analysis.

| **Sr. No.** | **Primary Antibody** |
| --- | --- |
| 1. | β-actin (1:1000, Mouse)  [ABclonal Technology, USA] |
| 2. | Insulin Receptor β (1:1000, rabbit)  [Cell Signaling Technology, USA] |
| 3. | IRS-1 (1:1000, Rabbit)  [ABclonal Technology, USA] |
| 4. | pIRS-1 Ser307 (1:1000, Rabbit)  [Cell Signaling Technology, USA] |
| 5. | Akt-1 (1:1000, Rabbit)  [ABclonal Technology, USA] |
| 6. | pAkt-1 S473 (1:1000, Rabbit)  [ABclonal Technology, USA] |
| 7. | Glut-4 (1:1000, Rabbit)  [ABclonal Technology, USA] |
| 8. | Anti-mouse IgG‑HRP (1:10000, goat)  [Genei, Bangalore] |
| 9. | Anti‑rabbit IgG‑HRP (1:5000, goat)  [Jackson ImmunoResearch, USA] |

**Table S3.** Antibodies used for the IHC studies.

| **Primary Antibody** | **Secondary Antibody** | **Excitation (nm)** | **Emission (nm)** |
| --- | --- | --- | --- |
| Anti-Insulin  (1:200, Guinea Pig)  [DAKO Agilent, USA] | Alexa 488 (1:500, Donkey)  [Jackson ImmunoResearch Laboratories, Inc. USA] | 493 | 519 |
| Anti-Glucagon (1:200,rabbit)  [Cell Signaling Technology, USA] | Alexa 647 (1:500, Donkey)  [Jackson ImmunoResearch Laboratories, Inc. USA] | 651 | 667 |
| BrdU (1:100, Rat)  [Abcam, USA] | Rhodamine Red (1:200, Donkey)  [Jackson ImmunoResearch Laboratories, Inc. USA] | 570 | 590 |
| NGN3 (1:50, rabbit)  [Thermo Fisher Scientific, USA] | Rhodamine Red (1:200, Rat)  [Jackson ImmunoResearch Laboratories, Inc. USA] | 570 | 590 |
| PDX-1 (1:1000, Goat)  [Abcam, USA] |  |  |  |
| ARX (1:500, Rabbit)  [Sigma-Aldrich, Germany] |  |  |  |
| PAX-4 (1:500, Goat)  [Sigma-Aldrich, Germany] |  |  |  |
| AIF (1:100, Rabbit)  [Abcam, USA] |  |  |  |
